# Pancreatic Schwann cell reprogramming supports cancer-associated neuronal remodeling

**DOI:** 10.1101/2023.12.20.572576

**Authors:** Martha M. Rangel-Sosa, Fanny Mann, Sophie Chauvet

**Affiliations:** Aix Marseille Univ, CNRS, IBDM, Marseille, France

**Keywords:** Pancreatic ductal adenocarcinoma, chronic pancreatitis, Schwann cells, axon sprouting, cell reprogramming, sympathetic nervous system, glial cell-derived neurotrophic factor

## Abstract

The peripheral nervous system is a key regulator of cancer progression. In pancreatic ductal adenocarcinoma (PDAC), the sympathetic branch of the autonomic nervous system inhibits cancer development. This inhibition is associated with extensive sympathetic nerve sprouting in early pancreatic cancer precursor lesions. However, the underlying mechanisms behind this process remain unclear. This study aimed to investigate the roles of pancreatic Schwann cells in the structural plasticity of sympathetic neurons. We examined the changes in the number and distribution of Schwann cells in a transgenic mouse model of PDAC and in a model of metaplastic pancreatic lesions induced by chronic inflammation. Schwann cells proliferated and expanded simultaneously with new sympathetic nerve sprouts in metaplastic/neoplastic pancreatic lesions. Sparse genetic labeling showed that individual Schwann cells in these lesions had a more elongated and branched structure than those under physiological conditions. Schwann cells overexpressed proinflammatory and neurotrophic factors, including glial cell-derived neurotrophic factor (GDNF). Sympathetic neurons upregulated the GDNF receptor and promoted cell growth in response to GDNF *in vitro*. Selective genetic deletion of *Gdnf* in Schwann cells completely blocked sympathetic nerve sprouting in metaplastic pancreatic lesions *in vivo*. This study demonstrated that pancreatic Schwann cells underwent adaptive reprogramming during early cancer development, supporting a protective antitumor neuronal response. These finding could help to develop new strategies to modulate cancer associated neural plasticity.

**MAIN POINTS:**

- Non-myelinating pancreatic Schwann cells associate with sympathetic axon terminals supplying the pancreas.
- Pancreatic Schwann cells proliferate and undergo adaptive reprogramming in response to chronic inflammation and the development of pancreatic cancer.
- Glial cell line-derived neurotrophic factor expression in reprogrammed pancreatic Schwann cells promotes Schwann cell expansion and sympathetic axon sprouting in pancreatic cancer precursor lesions.

## 1 INTRODUCTION

Schwann cells (SCs) are the main glial cells of the peripheral nervous system that nurture nerve fibers and provide vital metabolic support for axonal signaling. SCs remain quiescent under normal physiological conditions. When axonal contact is lost due to peripheral nerve injury, myelin and Remak SCs proliferate and undergo adaptive reprogramming, generating repair SCs that are specialized to promote axonal regeneration and reinnervation of distal targets (Jessen & Arthur-Farraj, 2019). This reprogramming involves important morphological changes as the cells adopt an elongated bipolar morphology and align to form axon guidance tracks (Büngner bands) (Jessen & Arthur-Farraj, 2019). The SC response to injury also requires the upregulation of the transcription factor c-Jun and c-Jun-dependent expression of glial cell-derived neurotrophic factor (GDNF), brain-derived neurotrophic factor (BDNF), and other neurotrophic factors. These factors support neurite outgrowth *in vitro* and promote regeneration when applied exogenously to injured nerves *in vivo* (Jessen & Mirsky, 2022). This observation suggests that SCs may support regeneration by releasing growth factors directly affecting neurons.

Although SCs have been extensively studied in the context of peripheral nerve injury, their roles in the sprouting of uninjured nerve terminals have received less attention. Early studies investigated the sprouting of motor muscle innervation induced by muscle paralysis or partial muscle denervation and showed that new axonal sprouts were consistently associated with processes extended by terminal Schwann cells (Duchen, 1971; Son & Thompson, 1995). In addition, SCs from severed peripheral nerves induce the sprouting of motor nerve terminals when transplanted into normal muscles (Son & Thompson, 1995). Despite these findings, whether SCs play an active role in promoting axonal sprouting in other contexts, such as inflammatory conditions and cancer development, remains unknown.

Significant axonal sprouting has been reported during the initiation of pancreatic ductal adenocarcinoma (PDAC) (Guillot et al., 2022; Sinha et al., 2017; Stopczynski et al., 2014). Sympathetic nerve fibers supplying the pancreatic acinar tissue branch out to form dense networks that innervate early noninvasive lesions of the pancreas, such as acinar-to-ductal metaplasia (ADM) and pancreatic intraepithelial neoplasia (PanIN). This local increase in sympathetic nerve terminals is believed to inhibit the progression of precursor lesions to PDAC (Guillot et al., 2022). However, the exact molecular and cellular mechanisms responsible for the sprouting of sympathetic nerve fibers during the initial stages of PDAC development remain unknown.

The abovementioned activities of SCs position them as potential candidates for directing neuronal remodeling during cancer development; this is in line with the presence of non-myelinating SCs in the PDAC tumor microenvironment and around non-invasive PanIN precursor lesions (Deborde et al., 2022; Demir et al., 2014; Sun et al., 2023; Xue et al., 2023). Cancer cells can induce phenotypic reprogramming of SCs, resembling injury-induced repair SCs (Deborde et al., 2016; Shurin et al., 2019). Tumor-associated SCs have been implicated as active contributors to cancer progression, influencing cancer cell migration and invasion and promoting a tumor-favorable immune and stromal response within the tumor microenvironment (Deborde & Wong, 2022). Despite these advances, whether SCs are associated with nerve fibers in tumors and their potential roles in promoting changes in neuronal innervation during cancer development remain to be explored.

This study aimed to investigate whether non-myelinating pancreatic Schwann cells (nm-pSCs) play a role in the cancer-induced remodeling of pancreatic sympathetic innervation. We used three-dimensional (3D) imaging of the mouse pancreatic tissue to visualize nm-pSCs and their interactions with sprouting sympathetic nerve fibers in a transgenic mouse model of PDAC. We analyzed the response of nm-pSCs to pancreatic injury in a mouse model of chronic pancreatitis (CP) that developed highly sympathetic-innervated metaplastic ADM lesions. Our study provides insights into developing novel approaches to modulate cancer associated neural plasticity.

## 2 MATERIALS AND METHODS

All animal procedures were performed following the guidelines of the French Ministry of Agriculture (approval number F1305521) and were approved by the Ethics Committee for Animal Experimentation of Marseille – CEEA-014 (APAFIS#27289-2020091713522725 v4).

### 2.1 Mouse strains

Wild-type mice with a C57BL/6 background were obtained from Janvier Labs (Le Genest-Saint-Isle, France). KIC mutant mice (*Kras^LSL-^*^G12D/+^; *Cdkn2a (Ink4a/Arf) ^lox/lox^; Pdx1-Cre*) were generated in-house by intercrossing *Kras^LSL-^*^G12D/+^ (Jackson et al., 2001), *Pdx1-Cre Cre* (G. Gu, Dubauskaite, & Melton, 2002), and *Cdkn2a^lox/lox^* (Aguirre et al., 2003) mice. *Sox10-iCreER^T2^* mice (*CBA;B6-Tg(Sox10-icre/ERT2)388Wdr/J*) were purchased from Jackson Laboratory (Sacramento, CA) (McKenzie et al., 2014). *R26R-EYFP* mice (*B6.129X1-Gt(ROSA)26Sortm1(EYFP)Cos/J*) (Srinivas et al., 2001), *S100b-EGFP* mice (*Tg(S100b-EGFP)^11Lgrv^)* (Vives, Alonso, Solal, Joubert, & Legraverend, 2003), *R26R-DTA* mice (*Gt(ROSA)26Sor^tm1(^DTA)Riet*) (Brockschnieder, Pechmann, Sonnenberg-Riethmacher, & Riethmacher, 2006), *Gdnf^lacZ^* mice (*Gdnf^tm1Rosl^*) (Moore et al., 1996) and *Gdnf^lox/lox^* mice (*Gdnf ^tm1.1Joao^*) (Kopra et al., 2015) were kindly donated. *Sox10-iCreER^T2^*mice were crossed with *R26R-EYFP*, *R26R-DTA*, or *Gdnf^lox/lox^*. The offspring of these crosses were named *Sox10- iCreER^T2^*; *R26R-EYFP, Sox10-iCreER^T2^*; *R26R-DTA*, or *Sox10-iCreER^T2^*; *Gdnf^Lox/Lox^*. Animals were housed in a specific pathogen-free facility in individually ventilated cages under controlled conditions (12 h light/dark cycle; humidity: 45–65%; room temperature (RT): 23 ± 2 °C) and were provided ad libitum access to water and food.

### 2.2 Tissue collection

Mice were anesthetized by an intraperitoneal (IP) injection of 100 mg/kg ketamine (Imalgene; Merial) and 10 mg/kg xylazine (Rompun; Bayer) and were intracardially perfused with 20 mL of cold phosphate-buffered saline (PBS, Cat. #: 16431755, Thermo Scientific), followed by 30 mL of cold 4% paraformaldehyde (PFA) (Cat. #: 15714-S; Electron Microscopy Sciences) in PBS. The digestive system was collected and post-fixed overnight (O/N) in 4% PFA at 4 °C. The pancreas was dissected and used for subsequent experiments.

### 2.3 Chronic pancreatitis induction

Chronic pancreatitis (CP) was induced in 7–10-week-old C57BL/6 or transgenic mice by hourly IP injections of cerulein (50 μg/kg; Cat. #: C9026 Sigma-Aldrich) in PBS. Mice were injected with PBS (vehicle) or cerulein eight times daily, twice weekly, for 2, 4, or 6 weeks. Three days after the final day of injection, the mice were intracardially perfused, and the pancreas was collected as described above. The animals were randomly assigned to treatment groups (cerulein versus vehicle). Male and female mice were included in this study because they exhibited similar responses to cerulein-induced CP injury and recovery (Obafemi et al., 2019). Animals injected with the vehicle for different durations showed no ADM or changes in neuronal components; therefore, they were pooled as controls. Cerulein-induced ADM is easily detectable; thus, the experimenter could not be blinded to the treatment during the analysis.

### 2.4 Immunostaining on tissue sections

Fixed pancreases were incubated O/N in 30% sucrose in PBS at 4 °C, included in optimal cutting temperature (OCT) compound (Cat. #: 361603E, VWR) and cut as 18-μm-thick cryostat serial sections. Slides were washed with PBS, permeabilized with 0.1% Triton X-100 (cat. #: 35501-15, Nacalai Tesque) in PBS, and blocked with blocking solution (2% donkey serum (DK; Cat. #: S2170, Dutscher), 2% Bovine Serum Albumin (BSA; Cat. #: 421501J, VWR), and 0.1% Triton X-100 in PBS) for 1 h at room temperature (RT). Primary antibody incubation was performed O/N at 4 °C in the same blocking solution. Slides were then washed with PBS and incubated with secondary antibodies for 2 h at RT. Nuclei were stained with DAPI (4’,6-diamidino-2-phenylindole, dihydrochloride) NucBlue (Cat. #: R37606; Thermo Fisher Scientific). Slides were mounted using Aqua-Poly/Mount (Cat. #: 18606-20, Polysciences). Antibodies used in this study are listed in supplementary (Supp.) Table 1.

### 2.5 Three-dimensional homemade printed chambers

The chamber that underwent 3D printing was designed using the computer-aided design software Catia V5. Stereolithography was performed with a Form 3 B printer and Black V4 Resin (Cat. #: RS-F2-GPBK-04, Formlabs), constructing the chamber layer by layer with a 100 µm thickness for each layer. Following the printing process, a 5-min immersion in Form Wash using isopropanol (Cat. #: 20922.320, VWR) was used as the solvent to remove the unpolymerized resin from the surface of the printed parts. Subsequently, the printed chamber underwent post-polymerization for 30 min at 60 °C in the Form Cure, utilizing a combination of thermal and UV curing. The supports used during the printing process were removed 48 h after curing to minimize the potential for partial deformation.

### 2.6 Immunostaining and clearing of whole-mount and thick sections of pancreas

Immunostaining and clearing were performed on whole-mount pancreas or thick sections according to the iDisco+ protocol (Guillot et al., 2022). For sectioning, the fixed pancreas was included in 4% low-melting agarose (cat. #: 16520-100, Invitrogen) and cut into 500-μm-thick slices using a vibratome (Vibratome VT 1000S, Leica). Sections were post-fixed for 1 h at RT in 0.1% PFA. Whole pancreas or thick sections were dehydrated in a methanol/PBS series (20%, 40%, 60%, 80%, 100% methanol, diluted in PBS; 20847.295, VWR), 1 h (whole pancreas) or 10 min (thick sections) each at RT, and then bleached in 5% hydrogen peroxide (H_2_O_2_; Cat. #: 216763, Sigma-Aldrich) diluted in methanol O/N at 4 °C. Samples were rehydrated in successive 1 h (whole pancreas) or 10 min (thick sections) baths of 100%, 80%, 60%, 40%, 20% methanol in PBS, permeabilized in 20% dimethylsufoxyde (DMSO; Cat. #: D/4121/PB08; Fisher Chemical), 0.16% TritonX-100, and 18.4 g/L of glycine (cat. #: 0167, VWR) in PBS for 1 day (whole pancreas) or 2 h (thick sections) at 37 °C, and then incubated with blocking solution PTWH (0.2% Tween-20 (Cat. #: 28829.183, VWR) and 10 mg/L heparin (cat. #: H7405, Sigma-Aldrich) in PBS) for 2 days (whole pancreas) or O/N (thick sections) at 37 °C. Tissues were incubated with primary antibodies diluted in PTWH, 5% DMSO, 3% DK serum for 7 (whole sections) or 1 (thick pancreas) days at 37 °C, washed several times with PTWH and incubated with secondary antibodies for 2–3 days (whole pancreas) or 5 h (thick sections) at 37 °C. Tissues were dehydrated with methanol/PBS and incubated O/N at RT in 66% dichloromethane (DCM; cat. #: 270947; Sigma-Aldrich) and 33% methanol. Finally, samples were immersed in 100% DCM for 20 min and kept in dibenzyl ether (DBE; cat. #: 305197; Sigma-Aldrich) to homogenize the refractive index during image acquisition. The thick sections were rinsed and kept in ethyl cinnamate (ECI; cat. #: W243000, Sigma-Aldrich). Before image acquisition, sections were mounted in 3D homemade printed chambers covered with coverslips and sealed with a melted paraffin–vaseline mixture (∼2:1 ratio) (Cat. #: P3558, Sigma-Aldrich. Vaseline pure, Mercurochrome Laboratories).

### 2.7 EdU assay

Mice treated with cerulein or vehicle received an IP injection of EdU (15 mg/kg) (cat. #: BCN-001-500; BaseClick) diluted in PBS two times weekly for the duration of cerulein or vehicle injections. The 18-μm-thick cryostat tissue sections were incubated for 30 min in EdU reaction buffer (0.1 M Tris-HCl pH 7.0 (Cat. #: 648317, Merck), 4 mM CuSO4 copper (II) sulfate (Cat. #: C3036; Sigma-Aldrich), 0.001 mM cyanine 3 azide (cat. #: BCFA-080-1, Sigma-Aldrich), and 0.1 M ascorbic acid (Cat. #: A4403; Sigma-Aldrich) diluted in water) to reveal EdU incorporation.

### 2.8 Tamoxifen injection

Tamoxifen (Cat. #: T5648-1G, Sigma) was dissolved in sunflower seed oil (cat. #: S5007, Sigma-Aldrich) at 15 mg/ml. *Sox10-iCreER^T2^;R26R-EYFP* mice received one IP injection at a dose of 5 mg/kg to induce sparse yellow fluorescent protein (YFP) labelling of nm-pSCs. For experiments requiring near-complete Cre-mediated recombination in nm-pSCs, *Sox10-iCreER^T2^;R26R-EYFP*, *Sox10-iCreER^T2^;R26R-DTA*, *Sox10-iCreER^T2^;Gdnf^lox/lox^* mice, or their control genotypes were administered IP injections at a dose of 100 mg/kg once a week for 3 weeks. Please refer to the figures for a detailed timeline of each experiment.

### 2.9 FACS sorting and qPCR

*S100b-EGFP* mice were treated with cerulein or vehicle for 4 weeks. On the day of FACS sorting, mice were intracardially perfused with PBS, and pancreata were immediately dissociated for 30 min in an OctoMACS (Miltenyi Biotec) using 10 mg/ml Collagenase II (Cat #: LS0004176, Worthington) and 1.5 mg/ml of DNAse I (Cat. #: DN25, Sigma) in 5 ml Hanks’ Balanced Salt Solution (HBSS) with Calcium and Magnesium (Cat. #: 24020117; Thermo Fisher Scientific). The following solutions were prepared using the HBSS calcium- and magnesium-free solution (Cat. #: 14175095; Thermo Fisher Scientific). The enzymatic reaction was stopped with 10 mM EDTA (Cat. #: 15575020, Invitrogen). The debris was removed from the preparation using a debris removal solution (cat. #: 130-109-398, Miltenyi Biotec) according to the manufacturer’s instructions. Cells were incubated for 15 min with an anti-CD16/32 antibody (clone 2.4G2) to block Fc receptors and then stained for 20 min using the following antibodies: CD45-BUV395, EpCAM-PeCy7, Thy1.2-APC, NGFR-PE, Sca1-APCC7, and Ly006C-V605. The CD45^+^, EpCAM^+^, Thy1.2^+^, Sca1^+^, and Ly006C^+^ cells were depleted using Dynabead anti-rat (Cat. #: 11035, Invitrogen Life Technologies) according to the manufacturer’s protocol. DAPI (Cat. #: 564907; BD Pharmingen) was used for live/dead staining. This gating strategy allowed for the depletion of epithelial cells (EpCAM^+^) and fibroblasts (Thy1-2^+^, Sca1^+^, and Ly006C^+^), which were also GFP^+^ in *S100b-EGFP* mice. Live GFP^+^ and GFP^+^NGFR^+^ cells (staining negative for other markers) were sorted as SCs. Three independent experiments were conducted. RNA was extracted using the TRIzol reagent (Cat. #: 15596018, Invitrogen), and cDNA was synthesized using SuperScript™ VI First-Strand Synthesis kit (Cat. #: 18080051; Thermo Fisher Scientific) following the manufacturer’s instructions. qPCR analysis using StepOnePlus Real-Time PCR Systems (Thermofisher) and PowerUp SYBR™ Green Master mix (cat. #: A25742; Applied Biosystems) was performed to corroborate the SCs identity using primers for SRY-Box Transcription Factor 10 (*Sox10)*, S100 Calcium Binding Protein B (*S100b)*, nerve growth factor receptor (*Ngfr)*, and GDNF Family Receptor Alpha 3 (*GFRa3)* transcripts. The mRNA expression of SCs reprogramming markers, cytokines, and neurotrophic factors was assessed using qPCR (see genes and primers in Supp. Table 2).

### 2.10 Sympathetic neuron culture

Coeliac superior mesenteric ganglia (CSMG) were isolated from adult *S100b-GFP* mice treated with cerulein or vehicle. Dissection was performed using a binocular loop, and the tissues were kept in cold dissection medium (HBSS-glucose containing 5 mM HEPES (Cat. #: 14175095; Thermo Fisher Scientific; cat. #: 15630, Gibco), 10 mM d-glucose (cat. #: G/0500/60, Fisher Chemical), and 100 U/ml Penicillin-Streptomycin (Cat. #: 15140-122, Gibco)), with pH adjusted to 7.5 with NaOH (Cat. #: 369704; Carlo Erba) throughout the procedure. After dissection, tissues were placed in a digestion solution containing HBSS-glucose and 5 mM CaCl_2_ (Cat. #: 1564A930, Fischer Scientific) and a combination of collagenase II (2 mg/ml, Cat. #: 17101-015, Life Technologies) and dispase (5 mg/ml; Cat. #: 17105-041, Life tech invitro). The tissues were incubated for 30 min at 37 °C without agitation. After the first incubation, the medium was removed, and fresh digestion solution was added for a second 30-min incubation. The tissues were then centrifuged at 1300 ×*g* for 10 min. The supernatants were removed, and the tissues were washed with L15 complete medium (L-15 medium) (Cat. #: 11415-049, Gibco), 5% fetal calf serum (Cat. #: A5256801, Gibco), and 100 U/ml Penicillin-Streptomycin. The tissues were then dissociated using Pasteur pipettes of different diameters, gently pipetting up and down approximately 10–15 times per pipetting while avoiding air bubbles. The dissociated tissues were then layered onto Percoll (Cat. #: 17089101, Cytiva) gradient in a 15-ml falcon tube, with 4 ml of 12.5% Percoll solution layered over 28% Percoll in L-15 medium. The tubes were then centrifuged for 20 min at 1300 ×*g*, resulting in cell pellets and debris remaining in the suspension. The supernatants were removed, and fresh L-15 medium was added. Another 5 min of centrifugation at 900 ×*g* was performed, and the cell pellets were resuspended in NB complete medium (Neurobasal A medium (Cat. #: 21103-049, Gibco), B27 (Cat. #: 12587-010, Gibco), 2 mM l-glutamine (Cat. #: 25030024; Life Technologies, Carlsbad, CA, USA), and 100 U/ml Penicillin-Streptomycin). Finally, the neurons were plated in 4-well plates coated with poly-L-lysine (1 mg/ml) (Cat. #: P2636-500MG, Sigma-Aldrich) and homemade laminin (Chauvet et al., 2007) in complete NB medium supplemented with growth factors [50 ng/ml Beta-NGF (Beta-nerve growth factor, Cat. #: N-240, Alomon Labs), 100 ng/ml GDNF (Cat. #: PHC7044, Life technologies)]. Cells were incubated for 48 h at 37 °C and 5% CO_2_.

Cells were fixed with 4% PFA and 30% sucrose in PBS for 30 min, washed, and incubated in 50 mM NH_4_Cl (Cat. #: AA0235, VWR) in PBS for 15 min for autofluorescence quenching. Permeabilization was performed with 0.1% Triton X-100 in PBS for 15 min. Cells were blocked in 2% BSA, 2% goat serum, 2% DK, and 0.1% Triton X-100 in PBS for 1 h and then incubated O/N with primary antibodies in the blocking solution. After washing, the cells were incubated with secondary antibodies diluted in the blocking solution for 1 h. Finally, the nuclei were stained with DAPI, and the slides were mounted using Aquapolymount (Cat. #: 18606-20, Polysciences).

### 2.11 RNAscope in situ hybridization

Sections of 12-µm-thickness from control and cerulein-treated mouse CSMG were used to detect mRNA expression using the RNAscope v2 kit (Cat. #: 323110, ACD-Biotechne), following the manufacturer’s instructions. Briefly, slides were air dried, incubated with H_2_O_2_ for 10 min, and exposed to the target retrieval solution at 98 °C for 5 min. Slides were cooled down in fresh distilled water, incubated for 3 min in 100% ethanol, and left to dry at RT. The next day, slides were treated with Protease III for 15 min at 40 °C and then incubated with the indicated probes mix (Supp. Table 3) after 2 h at 40 °C. Subsequently, signal amplification and fluorophore detection were performed according to the manufacturer’s instructions. Nuclei were stained with DAPI, and the slides were mounted using Aquapolymount.

### 2.12 Imaging acquisition and processing

Cryostat sections were imaged using an M2 apotome and a confocal fluorescence microscope (Zeiss LSM 780 or LSM 880) with a 20x objective.

Whole-cleared pancreatic samples were imaged using a light-sheet fluorescence microscope (UltraMicroscope Blaze, Miltenyi Biotec, Bergisch Gladbach, Germany) equipped with a 5.5 Megapixel sCMOS camera. We used a 4x/0.35 objective with a 17-mm working distance dipping cap. The system was operated using the Inspector microscope control software, version 5.1.328. Cleared thick sections were imaged using a confocal microscope (LSM 780) with a long 25x objective. The cultured neurons were imaged using an inverted fluorescence microscope (Zeiss AxioObserver D1) equipped with a 10× Plan-NEOFLUAR objective.

### 2.13 Data analysis and statistics

Mouse sample sizes were calculated by power analysis using the statistical approach available at https://biostatgv. sentiweb. fr. Previous studies (Gomez-Sanchez et al., 2017; Guillot et al., 2022; Ledda, Paratcha, & Ibañez, 2002; Shurin et al., 2019) or pilot experiments have provided guidance for determining the effect size. The statistical power was set at 90% and the significance level at 5%. No samples or animals were excluded from the analyses. All investigators were blinded to the genotype throughout data collection and analysis. Each experiment was repeated at least three times, and the data were consistently reproduced for each replicate. Statistical analyses were performed using GraphPad Prism version 9 (GraphPad Software, Inc.). Normal distribution of the data was assessed using the D’Agostino–Pearson omnibus, Shapiro–Wilk, and Kolmogorov–Smirnov tests. Based on the normality results, the statistical tests included the Mann–Whitney test, Kruskal–Wallis test, and one-sample Wilcoxon test, all conducted as two-sided tests. Detailed descriptions of the statistical tests used in each experiment are provided in the figure legends.

For two-dimensional (2D) image analysis, the quantification of 2D images was performed using Fiji (Schindelin et al., 2012). Regions of interest (ROIs) were manually defined based on tissue autofluorescence and/or SRY-Box Transcription Factor 9 (SOX9) staining of pancreatic lesions to measure the density of sympathetic fibers and nm-pSCs. The areas of positive Tyrosine Hydroxylase (TH) or GFRA3 signals within these ROIs were quantified and normalized to the area of the ROIs. For sympathetic fiber association with nm-pSCs, the length of TH^+^ fibers associated with GFRA3^+^ cells was measured using the NeuronJ plug-in (Meijering et al., 2004) and expressed as a percentage of the total TH^+^ fiber length. Similarly, the length of the GFRA3^+^ nm-pSC network, with or without TH^+^ fibers, was measured and expressed as a percentage.

ADM formation in the cerulein-treated pancreatic samples was assessed by measuring the surface area of the ADM regions and expressed as a percentage of the total pancreatic tissue area. The percentage of proliferating cells in randomly selected ROIs was calculated as the number of EdU^+^/GFRA3^+^ or KI-67^+^/GFRA3^+^ cells relative to the total number of GFRA3^+^ cells. For RNAscope analysis, the percentage of *Th*^+^ cell bodies that showed a GDNF Family Receptor Alpha 1 (*Gfra1*) or GDNF Family Receptor Alpha 2 (*Gfra2*) signal and the number of dots per cell were counted.

For 3D image analysis, the 3D volumes were analyzed using the Imaris 64 software (version 9.2.1 and 9.3.0; Bitplane, Zurich, Switzerland). The stacked images were converted into Imaris files (.ims) using an Imaris File Converter. Individual YFP^+^ cells co-localizing with the GFRA3^+^ signal and a single SOX10^+^ nucleus were selected to analyze the SC morphology in thick sections. The morphology of these nm-pSCs was reconstructed manually using the “Filament tracer” tool. The following parameters were collected in the Imaris statistics tab: “dendrite length” (mean and sum values), defined as the mean branch length and the total length of the nm-pSCs; “number of terminal points” number of process extremities; “filament no. dendrite branch pts,” defined as the number of branch points of the nm-pSCs processes; “highest branch level,” which represents the maximum number of times that a process branch in each cell. The “Surface” tool was used to reconstruct 3D images of the nm-pSCs and measure their volumes.

For cell culture, the NeuronJ plugin (Meijering et al., 2004) was used to quantify the longest axon length in sympathetic neuron cultures.

For qPCR, the results of the qPCR were represented as mean Ct values or fold-change calculated using the ΔΔCt method (Schmittgen & Livak, 2008) with *Actin beta* as a housekeeping gene to compare relative gene expression between control and cerulein conditions.

## 3 RESULTS

### 3.1 SCs in the pancreas were non-myelinating and were associated with sympathetic nerve fibers supplying the exocrine pancreas

To reveal the presence of SCs in the pancreas, we first immunolabeled pancreas sections from wild-type adult mice using two pan-SC markers: anti-S100B, which delineates the cytoplasmic domain of the cells (Figure 1a), and anti-SOX10, which highlights the cell nuclei (Figure 1b). Consistent with earlier research findings (Sunami et al., 2001; Ushiki & Watanabe, 1997), all S100B^+^ SCs in the pancreatic tissue lacked myelin basic protein (MBP) (Figure 1a), classifying them as nm-pSCs. In contrast, MBP signals were detected exclusively within the nerve bundles outside the pancreas (Supp. Figure 1a). Consistently, a significant colocalization of S100B^+^ or SOX10^+^ cells with known markers of non-myelinating SCs, including GFRA3 (Figure 1a-b) and neural cell adhesion molecule (NCAM) (Supp. Figure 1b), was noted. GFRA3 was detected in all SCs identified by S100B or SOX10 staining (Figure 1a, b), as well as in cells expressing nerve growth factor receptor (NGFR) (Figure 1c) and glial fibrillary acidic protein (GFAP) (Supp. Figure 1c). Notably, GFRA3 showed specificity for nm-pSCs, with no staining observed for other cell types within the exocrine pancreas. Therefore, GFRA3 was used as a reliable marker for labeling nm-pSCs in this study.

**FIGURE 1.**
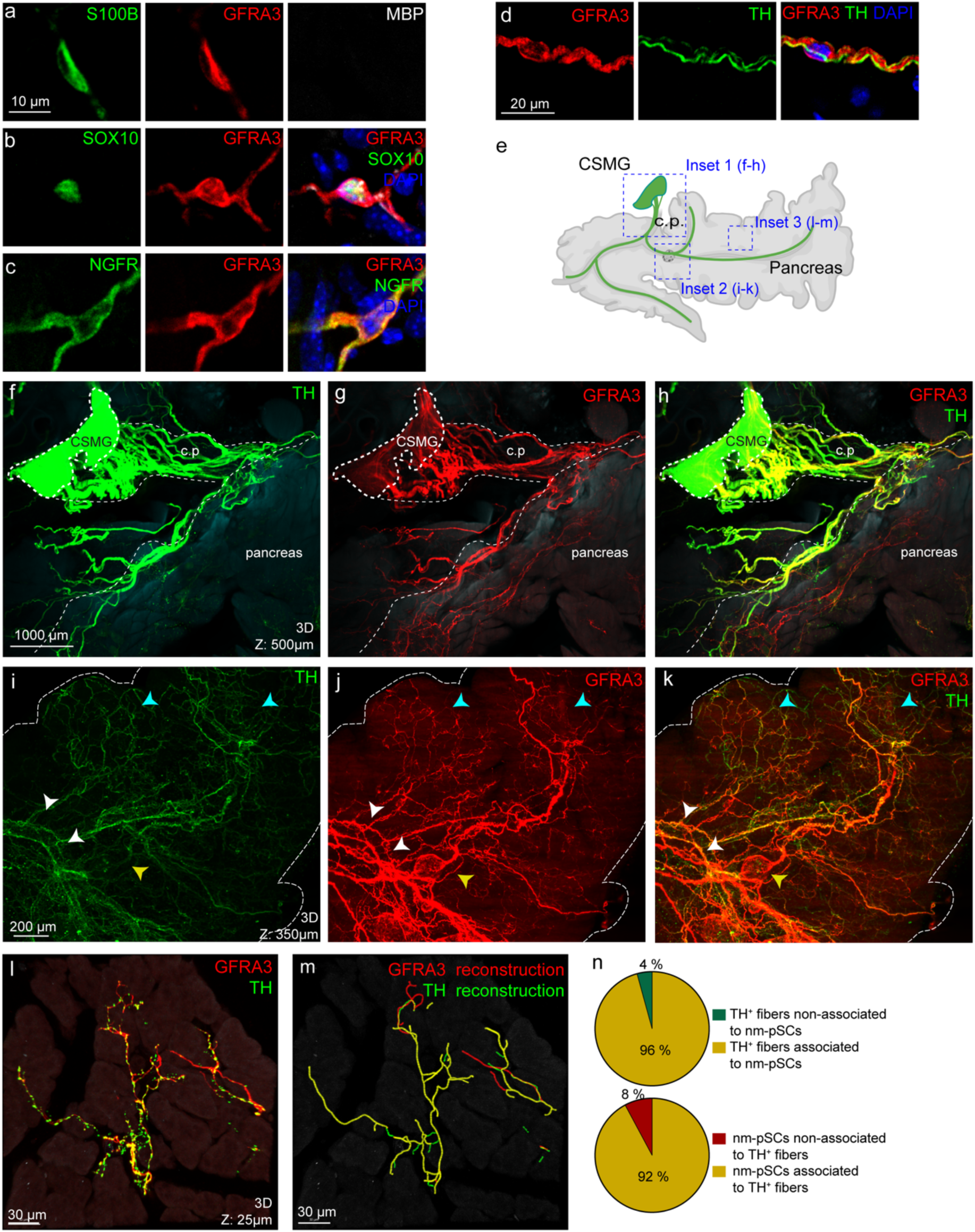
nm-pSCs associated with sympathetic nerve fibers in the mouse pancreas. (a–c) Representative images of cryostat sections of a mouse pancreas immunostained for the pan-SC markers S100B or SOX10, the myelin marker MBP, and the markers of nm-SCs GFRA3 and NGFR. Scale bar, 10 μm. (d) Representative images of cryostat sections of a mouse pancreas immunostained for GFRA3, TH, and DAPI. Scale bar, 20 μm. (e) Scheme illustrating the entire pancreas along with the CSMG complex, where the TH^+^ efferents (in green) extended towards the pancreas via the celiac plexus (c.p.). Upon penetrating the pancreas, the TH^+^ sympathetic nerves traverse alongside the principal pancreatic arteries. The insets 1, 2 and 3 provide reference for the spatial location of the pictures (f–h), (i–k), and (l–m), respectively. A single islet was depicted. (Made with biorender). (f–h) Maximum intensity projection of an optical section (500-μm thick) from a cleared mouse pancreas, coeliac plexus (c.p.), and adjacent CSMG double-stained for anti-GFRA3 and anti-TH antibodies. Scale bars, 1000 μm. (i–k) Maximum intensity projection of an optical section (350-μm thick) from a cleared mouse pancreas stained for anti-GFRA3 and anti-TH antibodies. nm-pSCs are observed surrounding the islets of Langerhans (yellow arrowhead), following TH^+^ nerve bundles (white arrowhead) and terminal TH^+^ fibers within the pancreas (blue arrowhead). Scale bars, 200 μm. (l, m) Maximum intensity projection of an optical section (25-μm thick) from a cleared mouse pancreatic lobule stained for anti-GFRA3 and anti-TH antibodies (l) and 3D reconstruction of GFRa3^+^ and TH^+^ signals (m). Scale bars, 30 μm. (n) Pie charts showing the percentage of the length of TH^+^ fibers associated with and without GFRA3^+^ nm-pSCs (n=3 mice, 23 images) and the percentage of GFRA3^+^ nm-pSCs signal associated with and without TH^+^ fibers (n=3 mice, 23 images). All images are representative observations in at least three mice. nm-pSCs, non-myelinating pancreatic Schwann cells; SOX10, SRY-Box Transcription Factor 10; S100B, Calcium Binding Protein B; MBP, myelin basic protein; GFRA3, GDNF Family Receptor Alpha 3; NGFR, nerve growth factor receptor; TH, Tyrosine Hydroxylase; CSMG, coeliac-superior mesenteric ganglion; c.p., coeliac plexus.

In pancreatic tissue sections, co-labeling with GFRA3 and TH revealed cellular-level colocalization, suggesting an association between nm-pSCs and TH^+^ sympathetic fibers (Figure 1d). We used 3D imaging to investigate the distribution of nm-pSCs and sympathetic nerve fibers throughout the pancreas. Sympathetic inputs to the pancreas originate from CSMG complex, forming large nerve bundles (coeliac plexus) that enter the pancreatic head and progressively defasciculate to innervate the entire organ, providing innervation to the islets and capillary bed of the exocrine tissue (Figure 1e) (Guillot et al., 2022; Rodriguez-Diaz et al., 2011). We observed that GFRA3^+^ nm-pSCs aligned with sympathetic fiber projections as soon as they emerged from the CSMG (Figure 1f–h); this association persisted on the intrapancreatic sympathetic nerve bundles (Figure 1i–k). In addition to the previously reported islet SCs at the pancreatic lobule level (Supp. Figure 1d), we also observed nm-pSCs covering the terminal sympathetic nerve fibers and extending through the acinar parenchyma (Figure 1l–m).

We quantified colocalization as a readout for the association between nm-pSCs and terminal sympathetic innervation in acinar tissue using 2D cryostat sections. Our results showed colocalization of GFRA3 signal exhibited along almost the entire length (96%) of TH^+^ sympathetic fibers. Conversely, approximately 8% of the GFRA3 signal was observed without any association with sympathetic fibers (Figure 1l-n). Instead, these nm-pSCs may associate with other nerve fiber types in the pancreas, such as cholinergic parasympathetic fibers (Supp. Figure 1e) or sensory fibers (Supp. Figure 1g–h).

These results showed that nm-pSCs formed an intricate network within healthy pancreatic tissue; almost all of them were superimposed on sympathetic nerves and terminal fibers that innervated the pancreas.

### 3.2 nm-pSCs covered sprouting sympathetic fibers during pancreatic cancer development

Next, we investigated the fate of sympathetic-associated nm-pSCs during PDAC development. We used the transgenic KIC (*Kras^LSL-^*^G12D/+^; *Cdkn2a (Ink4a/Arf) ^lox/lox^; Pdx1-Cre*) mouse model of PDAC at 6.5 weeks of age. At this stage, all mice developed at least one tumor, while earlier non-invasive precursor lesions could still be observed in the organ. KIC tissue samples were stained - either after sectioning or in wholemount - with a combination of antibodies targeting GFRA3, TH, and either SOX9 or cytokeratin 19 (CK19). The latter markers were used to identify pancreatic ductal cells, PDAC, and its precursor lesions. Consistent with previous findings (Guillot et al., 2022), we found an increased density of TH^+^ sympathetic fibers within ADM and PanIN lesions compared with that in control exocrine tissues (Figures 2a–c and 2e and Supp. Figure 2a). In contrast, PDAC tumors showed minimal presence of sympathetic fibers (Figures 2d and 2e and Supp. Figure 2a). As in the control pancreas, the GFRA3 antibody selectively labeled nm-pSCs in the pancreatic lesions of the KIC pancreas, as confirmed by their colocalizations with other SC markers, S100B, SOX10, NGFR, and GFAP (Supp. Figure 2b). An increase was observed in the area of GFRA3 immunostaining and in the GFRA3^+^ cell bodies in all types of pancreatic lesions, including ADM and PanIN precursor lesions, as well as in PDAC tumors (Figures 2a–d and 2f-g and Supp. Figure 2a). The density of TH^+^ fibers exhibited a robust positive correlation with the density of GFRA3^+^ nm-pSCs within the healthy tissue, ADMs and PanINs (Supp. Figure 2c), however, no significant correlation was observed in PDAC (Supp. Figure 2c). We then examined whether the association between mature sympathetic fibers and nm-pSCs described above was maintained during axonal sprouting. In ADM, PanIN, and PDAC lesions, the association was similar to that observed in healthy pancreas, with >90% of the length of the TH^+^ sympathetic network associated with GFRA3 signal (Figure 2h). Conversely, in early ADM and PanIN lesions, GFRA3^+^ nm-pSCs were largely found in association with TH^+^ fibers, as in the controls (Figure 2i). In contrast, in PDAC, a significant increase in the proportion of GFRA3^+^ signal was not associated with sympathetic fibers (Figure 2i), consistent with the observation that PDAC lacked substantial sympathetic innervation. This GFRA3^+^ signal could reveal the association of nm-pSCs with other types of fibers infiltrating the PDAC (Supp. Figure 2d).

**FIGURE 2.**
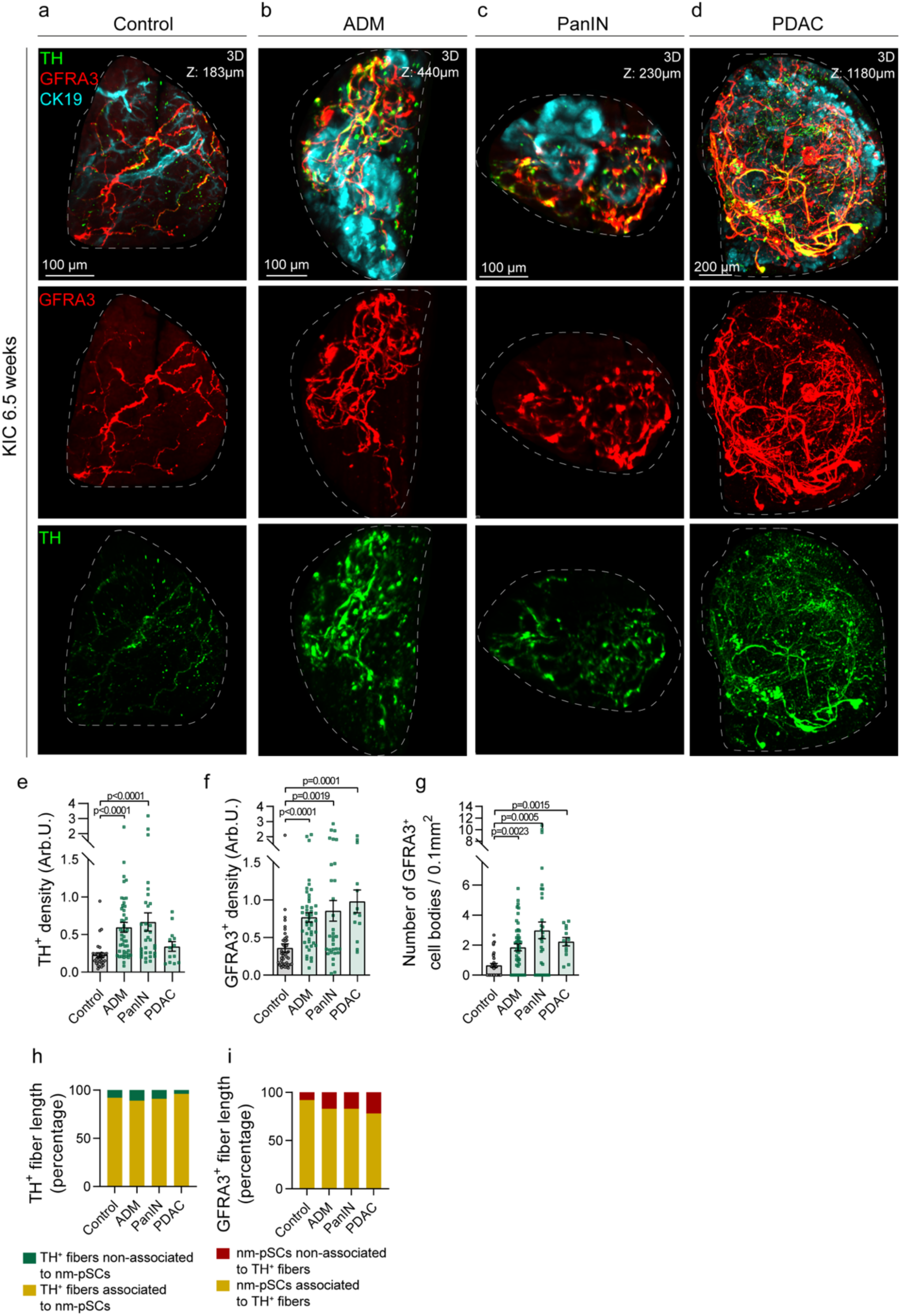
Coverage of sprouting sympathetic fibers by nm-pSCs during PDAC development. (a–d) Maximum intensity projection of optical sections (control (a): 183-μm thick, ADM (b): 440-μm thick, PanIN (c): 230-μm thick; PDAC (d): 1180-μm thick) through a cleared 6.5 week-old KIC pancreas stained with anti-GFRA3, anti-TH and anti-CK19 antibodies. The images are a representation of the observed staining in 3 different pancreata. Scale bars, 200 μm in PDAC and 100 μm in the other panels. (e–g) Scattered dot plots of TH^+^ signal density (e), GFRA3^+^ signal density (f), and GFRA3^+^ number of cell bodies (g) in acinar parenchyma of control mice and ADM, PanIN, and PDAC lesions of KIC mice. Number of mice: Control, 3; KIC, 3. Numbers of images analyzed: for (e), control, 32; ADM, 46; PanIN, 32; PDAC, 13; for (f and g), control, 41; ADM, 46; PanIN, 32; PDAC, 13. Data are presented as mean ± SEM. Statistical analysis: Kruskal–Wallis test with Dunn’s multiple comparisons, with the corresponding *p*-values shown in the figure. (h–i) Quantification of the percentage of the length of TH^+^ fibers associated or not associated with GFRA3^+^ nm-pSCs (h), and the percentage of GFRA3^+^ nm-pSCs signal associated or not associated with TH^+^ fibers (i). Numbers of mice: Control, 3; KIC, 3. Numbers of images analyzed: for (h and i), control, 18; ADM, 18; PanIN, 18; PDAC, 12. nm-pSCs, non-myelinating pancreatic Schwann cells; PDAC, pancreatic ductal adenocarcinoma; ADM, acinar-to-ductal metaplasia; PanIN, pancreatic intraepithelial neoplasia; KIC, *Kras^LSL-^*^G12D/+^; *Cdkn2a (Ink4a/Arf) ^lox/lox^; Pdx1-Cre*; GFRA3, GDNF Family Receptor Alpha 3; TH, Tyrosine Hydroxylase; CK19, cytokeratin 19.

These findings indicate that during the growth and branching of sympathetic axons in ADM and PanIN lesions, nm-pSCs expanded and covered the entire network of newly formed axonal branches.

### 3.3 Chronic pancreatitis induces expansion of sympathetic fibres and associated nm-pSCs in metaplastic lesions

A significant interaction between sympathetic axons and nm-pSCs during PDAC development suggests a potential role for nm-pSCs in nerve sprouting. However, studies on the underlying mechanisms in transgenic KIC mouse models are limited. Therefore, we established an alternative model to explore sympathetic axon remodeling induced by pancreatic injury. We used a cerulein-induced CP model to induce ADM lesions. Although sensory fiber remodeling has been shown in metaplastic lesions after 6 weeks of treatment in the CP model (Schwartz et al., 2013), potential changes in sympathetic innervation have not been described.

We subjected 8-week-old wild-type mice to CP treatment for 2, 4, and 6 weeks (Figure 3a) and monitored the evolution of metaplastic lesions and innervation in 2D sections and 3D-cleared pancreatic samples. ADM lesions, identified by the expression of ductal markers SOX9 and CK19, were visible after 2 weeks of treatment, and the proportion of metaplastic tissue in the pancreas progressively increased with the duration of the injections (Supp. Figure 3a– b). Similar to the ADM lesions in the KIC mouse model, we found a significant increase in the density of TH^+^ sympathetic fibers within the cerulein-induced ADM lesions at 2 weeks compared to the vehicle-injected tissue (Figures 3b–c and 3f). The density of TH^+^ sympathetic fibers did not change in histologically asymptomatic acinar tissues adjacent to the ADM (Figure 3f). From 2 to 6 weeks of treatment, high TH^+^ axon density was maintained within the ADM (Figure 3f). Since the ADM area increased with injection time (Supp. Figure 3c), the total number of TH^+^ sympathetic axons in the pancreas was expected to increase. We observed a simultaneous increase in the surface density of GFRA3 immunolabeling staining within ADM lesions (Figures 3b–e and 3g). Simultaneously, a progressive increase in the number of nm-pSCs was observed from 2 to 6 weeks, specifically within the ADM regions, with no changes in adjacent tissues (Figure 3h and Supp. Figure 3c and d). This increase in cell number was due to the proliferation of nm-pSCs, as demonstrated by EdU incorporation and Ki67 staining (Figure 3i–k and Supp. Figure 3e). Finally, we evaluated the association between nm-pSCs and sympathetic nerve fibers in ADM lesions. At all-time points analyzed, we observed that >90% of TH^+^ fibers were associated with the GFRA3 signal (Figure 3l). In contrast, as treatment progressed, an increasing percentage of GFRA3^+^ nm-pSCs without TH^+^ staining was observed at 6 weeks (Figure 3m). This GFRA3^+^ signal could reveal the association of nmSCs with sensory fibers infiltrating metaplastic lesions (Schwartz et al., 2013).

**FIGURE 3.**
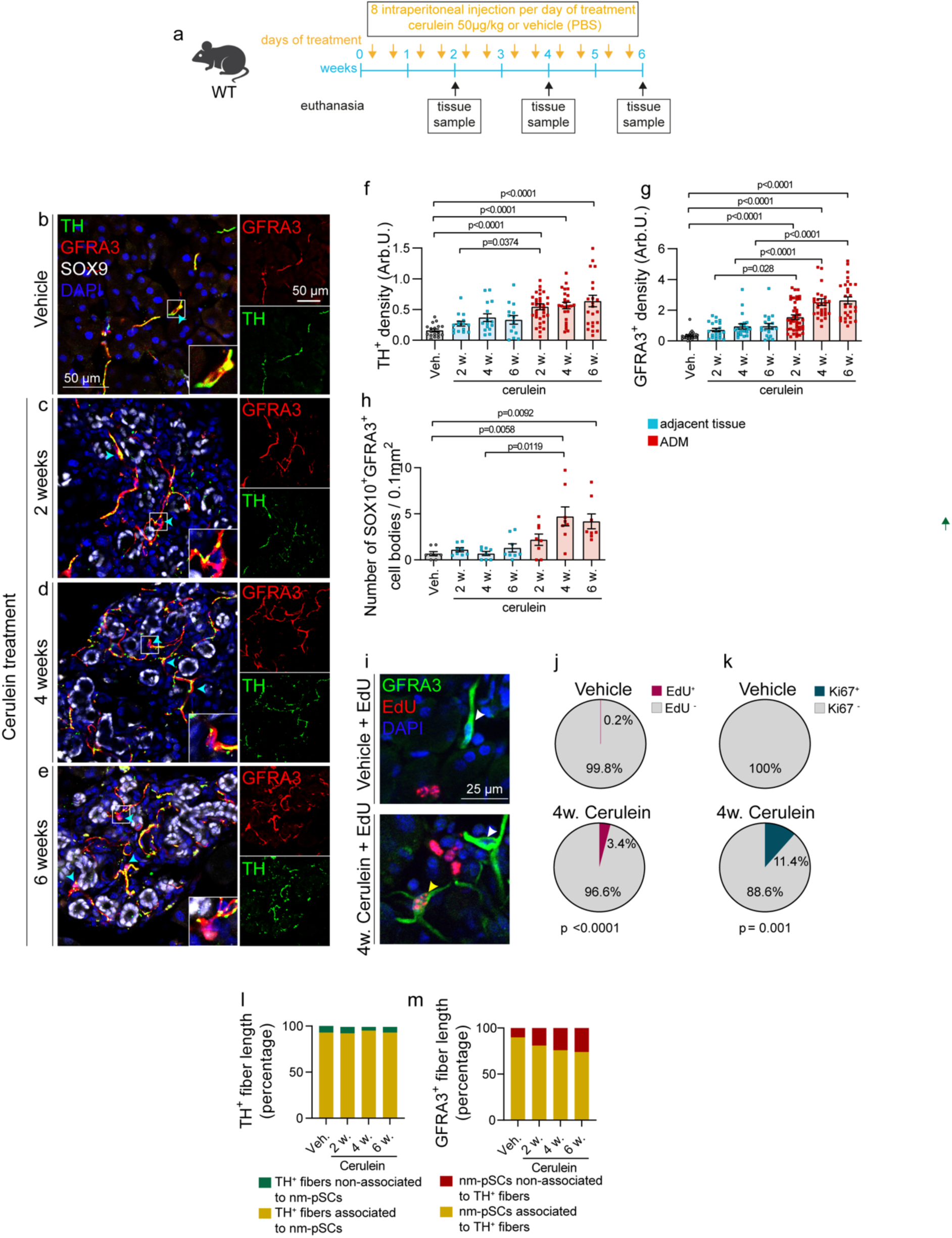
Coverage of sprouting sympathetic fibers by nm-pSCs in chronic pancreatitis. (a) Outline of induced-chronic pancreatitis experiment. (b–e) Representative images of pancreatic sections immunostained with anti-TH, anti-GFRA3 and anti-SOX9 antibodies showing sympathetic fibers and Schwann cells in acinar tissue injected with vehicle (b) or in ADM lesions induced by cerulein injections after 2 (c), 4 (d), or 6 (e) weeks of treatment. The cell bodies of the nm-pSCs were pointed with cyan arrowheads and one nm-pSCs was magnified. In each panel, separated channels of TH and GFRA3 staining were illustrated. Images are representative observations in at least three mice. Scale bars, 50 μm. (f–h) Scattered dot plots of TH^+^ signal density (f), GFRA3^+^ signal density (g), and number of GFRA3^+^SOX10^+^ cell bodies (h) in the pancreas of mice injected with vehicle (Veh.) for 6 weeks and in ADM and adjacent histologically asymptomatic tissues (adjacent tissue) in mice treated with cerulein for 2, 4, or 6 weeks. Numbers of mice: vehicle, 3; cerulein, 4. Numbers of images analyzed: for (f), vehicle, 18, cerulein/adjacent tissue, 15 (2 weeks), 15 (4 weeks), 14 (6 weeks), cerulein/ADM, 35 (2 weeks), 24 (4 weeks), 23 (6 weeks); for (g), vehicle, 24, cerulein/adjacent tissue, 21 (2 weeks), 21 (4 weeks), 20 (6 weeks), cerulein/ADM, 42 (2 weeks), 29 (4 weeks), 29 (6 weeks); for (h), eight pictures per condition were analyzed. Data are presented as mean ± SEM. Statistical analysis: Kruskal–Wallis test with Dunn’s multiple comparisons, with the corresponding *p*-values shown on the figure. (i) Representative images of EdU and GFRA3 staining on pancreatic cryostat sections from mice treated with vehicle or cerulein for 4 weeks. EdU^+^ (yellow arrowhead) and EdU^-^ (white arrowhead) nm-pSC are pointed. Scale bars, 25 μm. EdU was injected twice per week during vehicle or cerulein treatment. (j) Pie charts showing the percentages of EdU^+^ and EdU^-^ nm-pSCs in the pancreas of vehicle or cerulein treated mice at 4 weeks (see outline of EdU injections in Supp. Figure 3g). Vehicle: n=3 mice, one EdU^+^ cell out of 563 GFRA3^+^ cells; Cerulein: n=3 mice, 29 EdU^+^ cells out of 876 GFRA3^+^ cells. Statistical analysis used= Chi-square test, with the corresponding *p*-value shown in the figure. (k) Pie charts showing the percentages of Ki67^+^ and Ki67^-^ nm-pSCs in the pancreas of vehicle or cerulein-treated mice at 4 weeks. Vehicle: n=3 mice, 31 images analyzed, zero Ki67^+^ cell out of 92 GFRA3^+^ cells; Cerulein: n=3 mice, 27 images analyzed, 19 Ki67^+^ cells out of 166 GFRA3^+^ cells. Statistical analysis used the Chi-square test, with the corresponding *p*-value shown in the figure. (l–m) Quantification of the percentage of the length of TH^+^ fibers associated or not associated with GFRA3^+^ nm-pSCs (l) and the percentage of GFRA3^+^ nm-pSCs signal associated or not associated with TH^+^ fibers (m) in the pancreas of vehicle or cerulein treated mice. Numbers of mice: vehicle, 3; cerulein, 4. Numbers of images analyzed: Vehicle, 23, cerulein, 24 (2 weeks), 20 (4 weeks), 24 (6 weeks). nm-pSCs, non-myelinating pancreatic Schwann cells; TH, Tyrosine Hydroxylase; GFRA3, GDNF Family Receptor Alpha 3; SOX9, SRY-Box Transcription Factor 9; SOX10, SRY-Box Transcription Factor 10.

These findings suggest that CP induces changes in both neurons and glial cells that closely mimic those observed in early pancreatic lesions in KIC mice, making it a suitable model for investigating nm-pSCs in the context of an expanding sympathetic axonal network.

### 3.4 nm-pSC ablation in CP triggers compensatory mechanisms

To study the function of nm-pSCs in CP-induced sympathetic axon growth, we performed targeted ablation of nm-pSCs by crossing *R26R-DTA* mice, which conditionally expressed diphtheria toxin subunit A (DTA), with *Sox10-iCreER^T2^*mice. Mice were injected intraperitoneally with tamoxifen (100 mg/kg) once a week for a total of 3 weeks (Figure 4a) to achieve Cre recombination in as many cells as possible. In the reporter *Sox10-iCreER^T2^; R26R-EYFP* mice, this dosing protocol induced YFP expression in 81% of nm-pSCs (Suppl. Figure 4a–c). One week after the last tamoxifen injection, we quantified the GFRA3 immunoreactive signal in the pancreas of *Sox10-iCreER^T2^; R26R-DTA* mice. This analysis revealed an approximately 50% reduction in the density and number of nm-pSCs in healthy and adjacent tissue (Figure 4c and d and Supp. Figure 4d-e) and no noticeable effect on sympathetic axon density (Figures 4e and Supp. Figure 4d-e). Despite this reduction in nm-pSCs, when *Sox10-iCreER^T2^;R26R-DTA* mice were exposed to 2 weeks of cerulein treatment, both the surface density and number of GFRA3^+^ nm-pSCs increased selectively within the ADM lesions to levels comparable to *R26R-DTA* control mice (Figure 4b-d). Notably, we also observed a similarly high density of TH*^+^*sympathetic fibers in ADM lesions, regardless of whether the mice had undergone SC-targeted ablation of nm-pSCs (Figures 4b and 4e); this suggests that the nm-pSCs, regardless of their initial levels, can proliferate and adjust their numbers to match the increasing density of sympathetic axons precisely.

**FIGURE 4.**
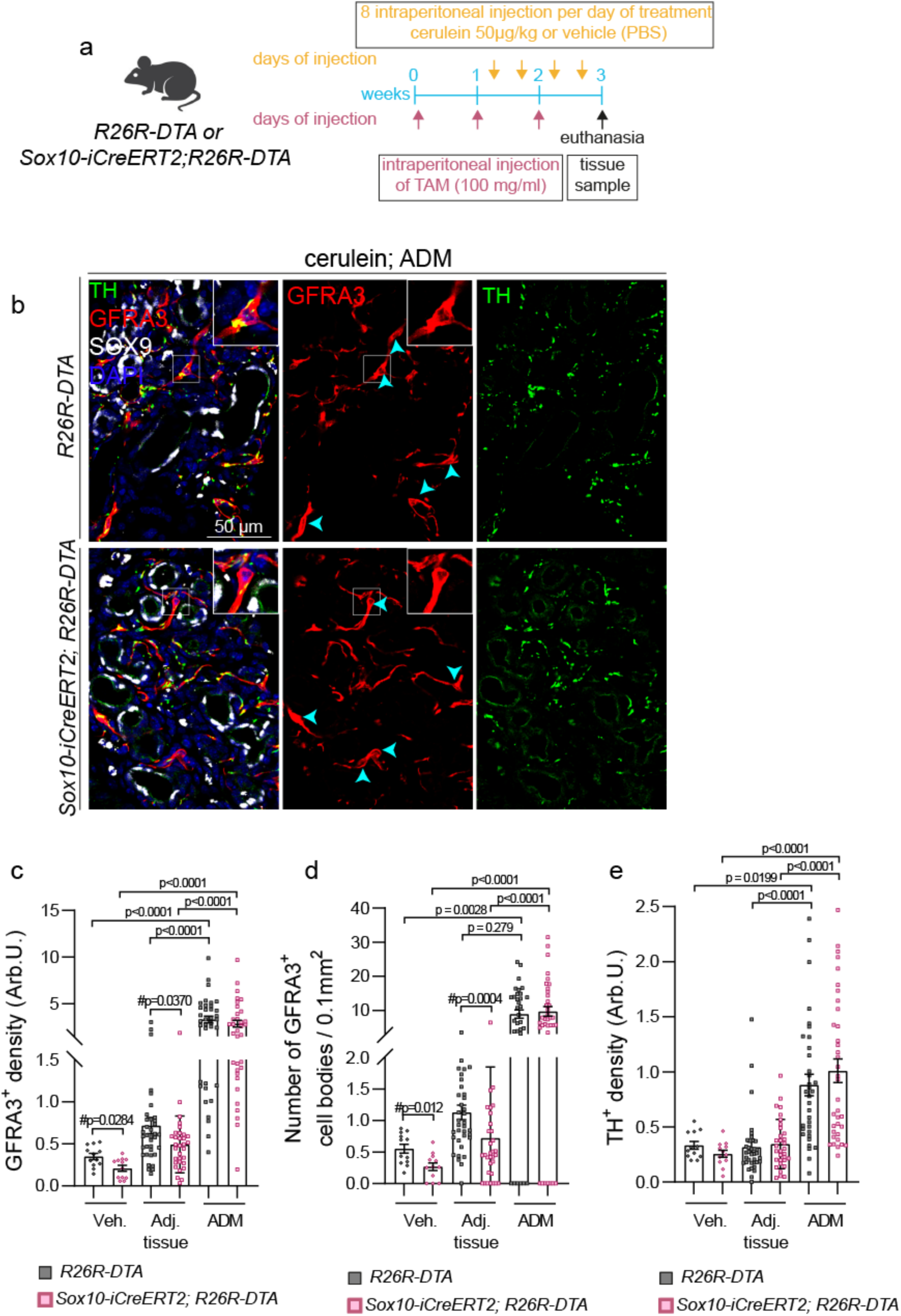
Effect of nm-pSCs ablation and compensatory mechanisms in chronic pancreatitis. (a) Outline of the experimental procedure for combined genetic depletion of nm-pSCs in *Sox10-iCreERT2;R26R-DTA* mice and 2-week induction of chronic pancreatitis. (b) Representative images of pancreatic cryostat sections from vehicle-injected *R26R-DTA* or *Sox10-iCreER^T2^;R26R-DTA* mice immunostained with anti-GFRA3 and anti-TH antibodies. Cell nuclei were counterstained with DAPI. nm-pSCs cell bodies were pointed with cyan arrowheads and one nm-pSCs was magnified. Images are representative observations in at least three mice. Scale bar,100 μm. (c–e) Scattered dot plots of TH^+^ signal density (c), GFRA3^+^ signal density (d), and the number of GFRA3^+^ cell bodies (e) in the pancreas of *R26R-DTA* or *Sox10-iCreERT2;R26R-DTA* mice treated with vehicle (Veh.) or cerulein for 3 weeks. In cerulein-treated mice, quantification was performed in the ADM and adjacent histologically asymptomatic tissue (Adj. tissue). Numbers of mice: *R26R-DTA*, 3 (vehicle) and 3 (cerulein); *Sox10-iCreERT2;R26R-DTA*, 3 (vehicle) and 3 (cerulein). Numbers of images analyzed for (e-g): *R26R-DTA*, 12 (veh.), 36 (adj. tissue), 34 (ADM); *Sox10-iCreERT2;R26R-DTA*, 12 (veh.), 32 (adj. tissue), 36 (ADM). Data are presented as mean ± SEM. Statistical analysis: Kruskal–Wallis test with Dunn’s multiple comparisons or Mann–Whitney test is indicated by #, with the corresponding *p*-values shown in the figure. nm-pSCs, non-myelinating pancreatic Schwann cells; TH, Tyrosine Hydroxylase; GFRA3, GDNF Family Receptor Alpha 3; ADM, acinar-to-ductal metaplasia; veh., vehicle; adj., adjacent; DTA, diphtheria toxin subunit A.

### 3.5 nm-pSCs underwent extensive morphological remodeling during metaplasia

Previous studies on SCs in injured nerves and terminal SCs in denervated muscles showed that SCs can extend processes to guide axon regrowth and sprouting (Gomez-Sanchez et al., 2017; Son & Thompson, 1995). Such a process extension mechanism could explain the observed increase in GFRA3 surface density in ADM lesions in parallel with nm-pSC proliferation. Therefore, we used a genetic strategy to label isolated nm-pSCs with YFP and analyzed their morphological changes during CP. We generated *Sox10-iCreER^T2^;R26R-EYFP* mice, in which a small number of *Sox10*-expressing cells were induced to express YFP by administration of a single low dose (5 mg/kg) of tamoxifen. One week later, the mice were treated with either vehicle or cerulein for 2 weeks (Figure 5a). Cleared thick sections of the pancreas were imaged in 3D printed chambers (Supp. Figure 5a) by confocal microscopy to visualize the isolated YFP-labelled nm-pSCs. Single-cell labeling was confirmed by anti-SOX10 nuclear staining (Figure 5b), and the SC identity of YFP^+^ cells was confirmed by GFRA3 co-staining (Figure 5b). The 3D morphology of individual SOX10^+^YFP^+^ nm-pSCs was then reconstructed and quantified using the Imaris software, following the pipeline described in Figure 5c.

**FIGURE 5.**
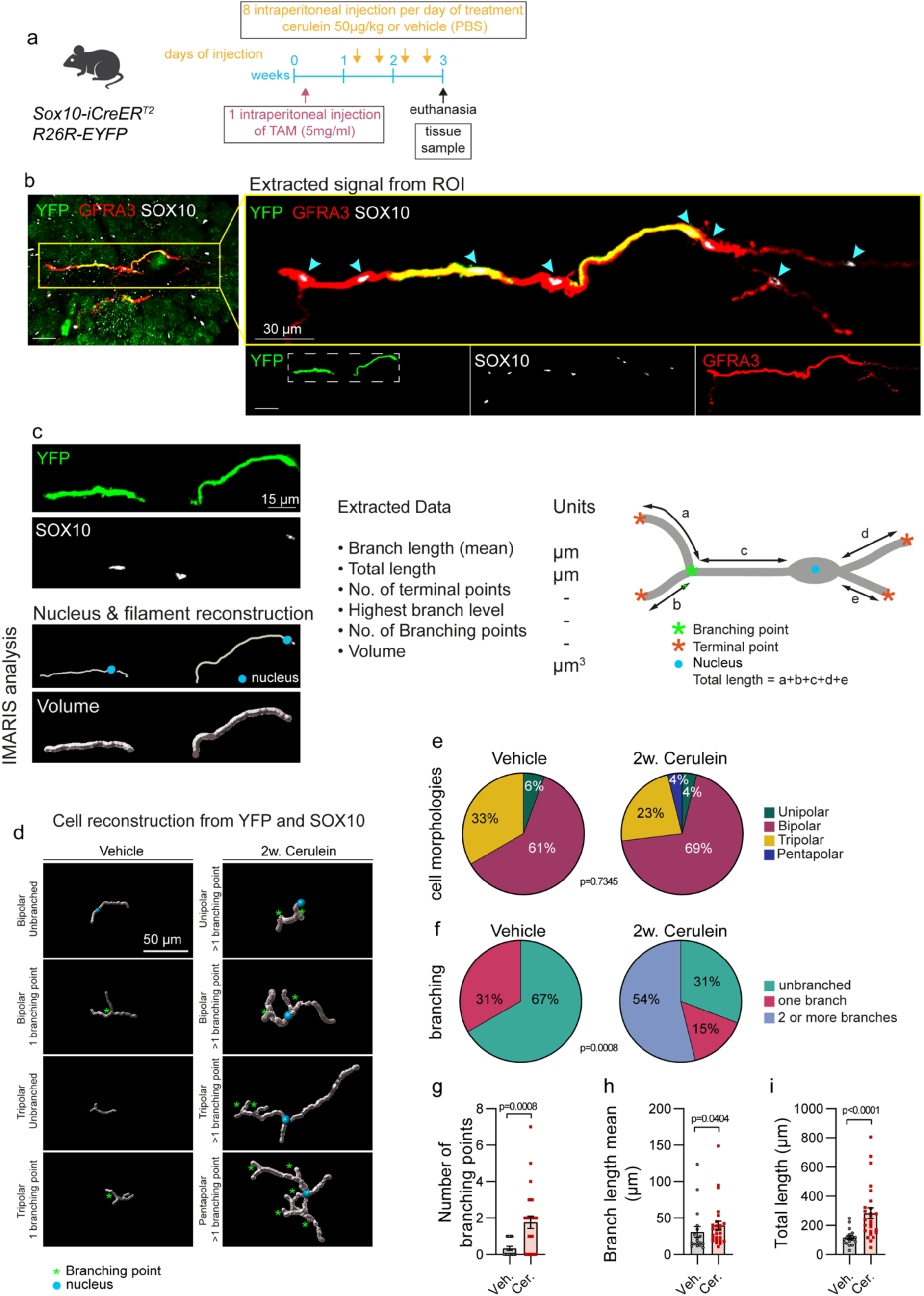
Morphological remodeling of nm-pSCs in metaplasia. (a) Outline of the experiment of combined nm-pSC sparse genetic labeling in *SOX10-iCreERT2;R26R-EYFP* mice and the 3-week induction of chronic pancreatitis. (b) Confocal image of a thick cleared section of *Sox10-iCreERT2;R26R-EYFP* mouse pancreas stained with anti-GFRA3, anti-YFP, and anti-SOX10 antibodies. Magnification of the signal extracted from the ROI is shown (blue arrowheads indicate SOX10^+^ nuclei). Individual channels for YFP, SOX10, and GFRA3 are also shown. Scale bars, 30 μm in the magnified triple staining and the individual stainings. (c-d) 3D reconstruction of individual nm-pSC morphologies from YFP and SOX10 fluorescence signals. (c) The quantified parameters are presented in detail in the figure. Scale bars, 15 μm. (d) Examples of reconstructed nm-pSC morphology observed in the pancreas of mice treated with vehicle or cerulein for 3 weeks. The blue circles represent the nucleus, the silver lines represent the processes, and the green asterisks represent branching points. Scale bars, 50 μm. (e) Pie chart showing the percentages of unipolar, bipolar, tripolar, and pentapolar nm-pSCs in the pancreas of vehicle- and cerulein-treated mice. Vehicle: n=3 mice, 18 nm pSCs; cerulein: n=3 mice, 26 nm pSCs. Statistical analysis was performed using the Chi-square test, and the corresponding *p*-values were shown. (f) Pie charts showing the percentage of unbranched nm-pSCs with one branch or with two or more branches in the pancreas of vehicle- or cerulein-treated mice. Vehicle: n=3 mice, 18 nm pSCs; cerulein: n=3 mice, 26 nm pSCs. Statistical analysis was performed using the Chi-square test, and the corresponding *p*-values were shown. (g–i) Quantification of branching points (g), mean branch length (h), and total length (i) of nm-pSCs in the pancreas of mice treated with vehicle or cerulein. Data are presented as mean ± SEM; Vehicle: n=3 mice, 18 nm pSCs; Cerulein: n=3 mice, 26 nm pSCs. Statistical analysis: Mann–Whitney test, with the corresponding *p*-values shown in the figure. nm-pSCs, non-myelinating pancreatic Schwann cells; SOX10, SRY-Box Transcription Factor 10; GFRA3, GDNF Family Receptor Alpha 3; YFP, yellow fluorescent protein; ROI, Regions of interest; Veh., vehicle; Cer., cerulein.

In both conditions, we observed three main categories of morphologies based on processes originating from the cell body: unipolar cells with a single process, bipolar cells with two processes, and tripolar cells with three processes emanating from the cell body (Figure 5d–e). Rare pentapolar cells were also observed (Figure 5d–e). Under control conditions, these primary processes typically remained unbranched or had only one branching point, as shown in Figures 5d and 5f, respectively. In the CP model, these branching morphologies represent only 46% of the total nm-pSC population. Instead, 54% of nm-pSCs showed an increase in process branching (Figure 5f), reflected in an increase in branching points (Figure 5g), branch length mean (Figure 5h), number of terminal points (Supp. Figure 4b), and the highest branch level per cell (Supp. Figure 5c). Consequently, the total length (Figure 5i) and volume (Supp. Figure 5d) of the nm-pSCs increased significantly in the CP group, consistent with the increased GFRA3 signal density observed in previous experiments.

These results showed that in parallel with the structural remodeling of sympathetic neurons, major changes occurred in the morphology of nm-pSCs during inflammation, leading to longer processes and increased branching complexity.

### 3.6 nm-pSCs underwent molecular reprogramming during metaplasia

The striking morphological changes observed in nm-pSCs during CP suggest that inflammation induces a cellular state similar to that observed in repair SCs following nerve injury (Jessen & Arthur-Farraj, 2019). This phenotypic transformation is likely associated with significant changes in gene expression and activation of the innate immune response, inducing cytokine upregulation. Here, we established a FACS sorting strategy to purify nm-pSCs from the pancreas of *S100b-EGFP* mice with or without CP (Supp. Figure 6) and analyzed the gene expression. We observed that nm-pSCs acquired a pro-inflammatory profile during CP, in particular with overexpression of interleukin-6 (*il-6*), transforming growth factor beta (*Tgfb*), C-C motif chemokine ligand 2 (*Ccl2*), interleukin-1 beta (*il-1b*), and tumor necrosis factor-alpha (*tnf*) (Figure 6a).

**FIGURE 6.**
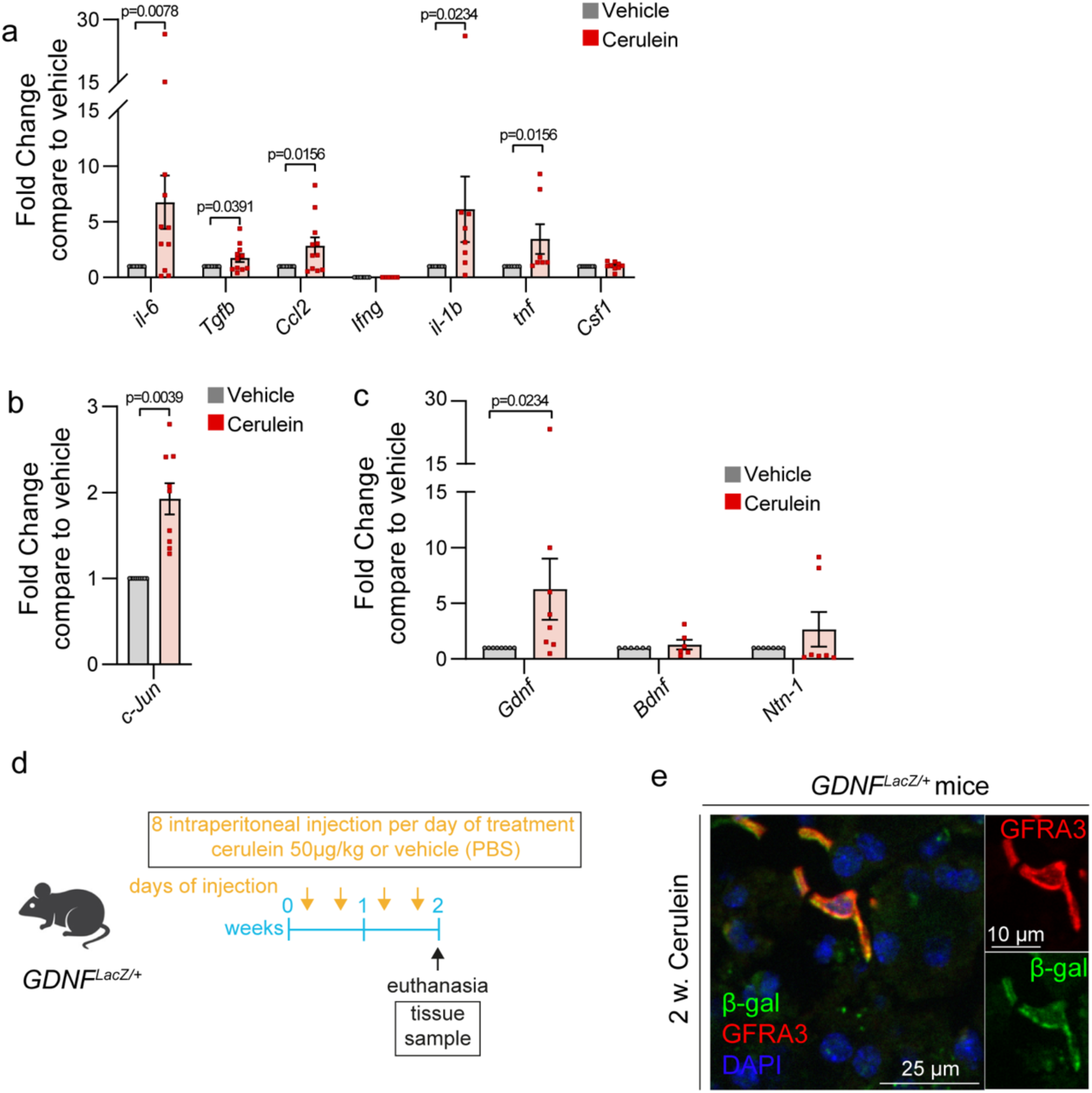
Molecular reprogramming of nm-pSCs in metaplasia. (a) qPCR analysis of mRNA expression of cytokines in pancreatic FACS-sorted nm-pSCs of mice treated with a vehicle or cerulein for 2 weeks. Each dot represents nm-pSCs from two pancreata. Fold change means compared to vehicle: *il-6*, 8.217; *Tgfb*, 2.00; *Ccl2*, 2.84; *ifng*, 0; *il-1b*, 6.121; *tnf*, 3.452; *Csf1*, 1.035; n= 6-10 samples tested in triplicate. Data are presented as mean ± SEM. Statistical analysis used in the One sample Wilcoxon test, with the corresponding *p*-values shown in the figure. (b) qPCR analysis of *c-Jun* mRNA expression of mice treated with vehicle or cerulein for 4 weeks. Data are presented as mean ± SEM. Fold change mean compared with vehicle: *c-Jun,* 1.927. Each dot represents nm-pSCs from two pancreata and tested in triplicate for vehicle or cerulein. Statistical analysis used the One sample Wilcoxon test, with the corresponding *p*-value shown in the figure. (c) qPCR analysis of mRNA expression of candidate neurotrophic factors and axon guidance cues in pancreatic FACS-sorted nm-pSCs of mice treated with a vehicle or cerulein for 2 weeks. Each dot represents nm-pSCs from two pancreata. Fold change means compared to the vehicle: *Gdnf*, 7.780; *Bdnf*, 1.288; *Ntn-1*, 2.654; n= 6–10 samples tested in triplicate. Data are presented as mean ± SEM. Statistical analysis used the One sample Wilcoxon test, with the corresponding *p*-values shown in the figure. (d) Outline of induced-chronic pancreatitis experiment in GdnflacZ/+ mice. (e) Representative images of pancreatic cryostat sections from 2 weeks (2 w.) cerulein-treated *Gdnf^lacZ^* mice, showing immunostaining with anti-beta-galactosidase (β-gal) and anti-GFRA3 antibodies. One nm-pSC was magnified to provide a clearer view of individual labeling. Images are representative observations in at least three mice. Scale bars, 25 μm and 10 μm in magnified panel. nm-pSCs, non-myelinating pancreatic Schwann cells; il-6, interleukin-6; Tgfb, transforming growth factor beta; Ccl2, C-C motif chemokine ligand 2; ifng, interferon gamma; il-1b, interleukin-1 beta; tnf, tumor necrosis factor-alpha; Csf1, colony stimulating factor 1; GFRA3, GDNF Family Receptor Alpha 3; GDNF, glial cell-derived neurotrophic factor. Bdnf, brain-derived neurotrophic factor; Ntn-1, Netrin-1.

In the context of nerve injury, transcription factor Jun proto-oncogene (c-Jun) has been extensively documented as the primary driver of SC reprogramming (Arthur-Farraj et al., 2012; Jessen & Mirsky, 2022). Using qPCR, we noted *c-Jun* mRNA overexpression in nm-pSCs from mice with CP compared to controls (Figure 6b). Next, using a candidate approach, we studied whether nm-pSCs upregulated factors known to be secreted by repair SCs to stimulate nerve regeneration and whose expression was upregulated by *c-Jun*. The expression of the neurotrophic gene *Bdnf* and the axon guidance gene netrin-1 (*Ntn1*) showed no significant increase in nm-pSCs between the control and CP groups (Figure 6c). In contrast, the neurotrophic gene *Gdnf* showed increased expression in nm-pSCs during CP (Figure 6c). We confirmed GDNF expression *in vivo* using *Gdnf^lacZ^* reporter mice, which showed beta-galactosidase staining in GFRA3^+^ nm-pSCs in the pancreas subjected to CP for 2 weeks (Figures 6d and 6e). Thus, pancreatic inflammation induced the molecular reprogramming of nm-pSCs, suggesting the role of nm-pSCs in initiating or amplifying the inflammatory response and supporting axon regrowth.

### 3.7 GDNF expression by nm-pSCs was required for SC expansion and sympathetic axon sprouting

As GDNF can induce neurite outgrowth from embryonic sympathetic ganglia (Ledda et al., 2002), we investigated its role in adult sympathetic neurons. GDNF functions primarily through the GFRA1 receptor (Klein et al., 1997; Sanicola et al., 1997); although it can also use GFRA2 to some extent (Airaksinen & Saarma, 2002). We first confirmed the expression of these two receptors in adult sympathetic neurons of the CSMG using RNAscope *in situ* hybridization (Figure 7a). Quantification of the percentage of neuronal cell bodies coexpressing *Th* and *Gfra1* mRNAs and the number of *Gfra1* mRNA signal dots per neuron revealed a significant increase in gene expression after cerulein treatment (Figure 7a–c). A similar observation was made for *Gfra2* mRNA (Figures 7a and 7d–e). *Gfra1* and *Gfra2* mRNA expressions were also detected in the FACS-isolated nm-pSCs, but the levels remained unchanged between the control and CP conditions (Figure 7f).

**FIGURE 7.**
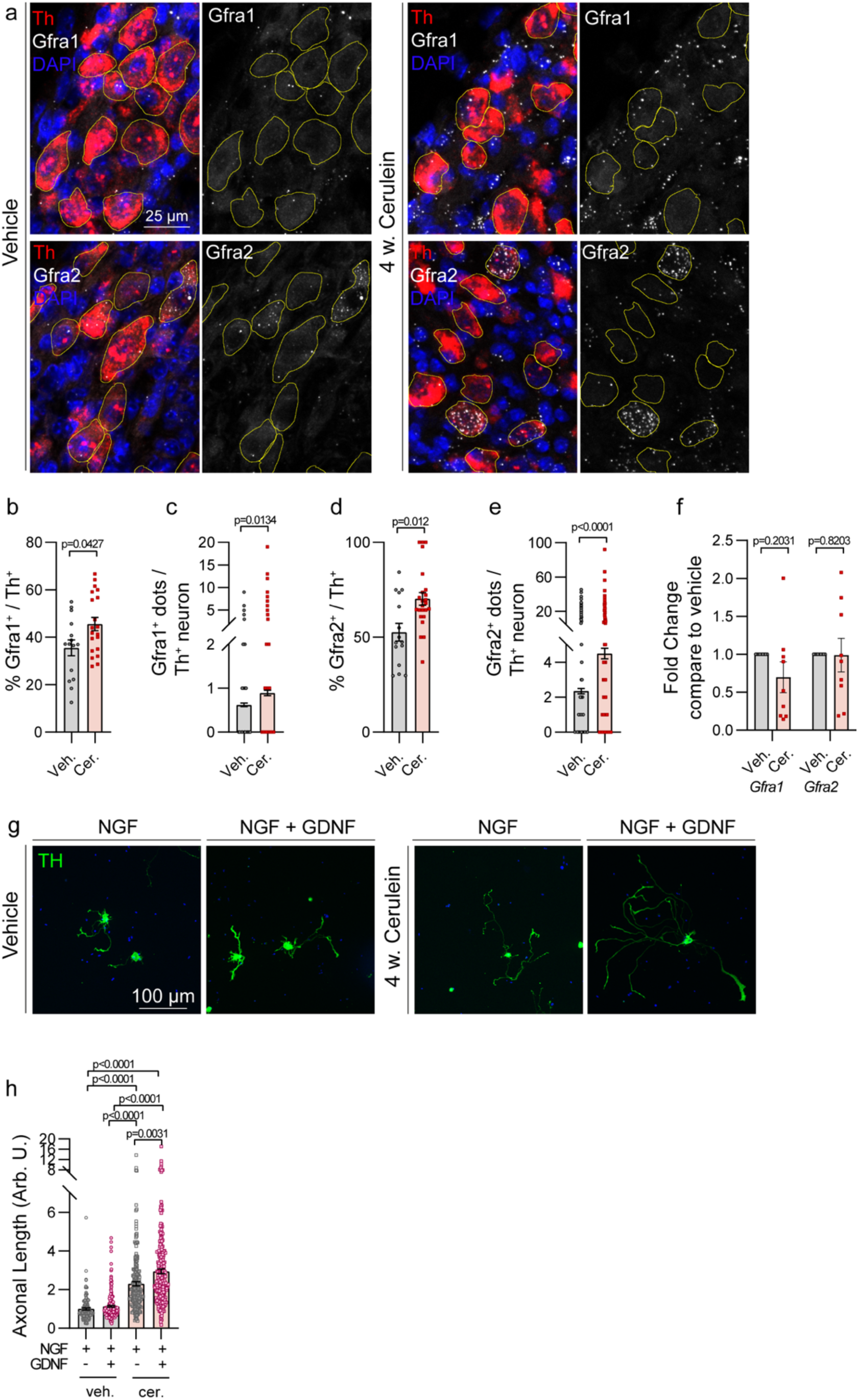
Sympathetic neurons expressed GDNF receptors and responded to GDNF *in vitro*. (a) Representative images of CSMG sections from mice treated with vehicle or cerulein for 4 weeks showing RNAscope signals for *Th* together with *Gfra1* or *Gfra2*. Yellow lines outline sympathetic neuronal cell bodies based on *Th* staining. Scale bar, 25 μm. (b–e) Scatter plots depicting the percentage of *Th^+^* neurons expressing *Gfra1* (b) or *Gfra2* (c), as well as the number of *Gfra1^+^*(d) or *Gfra2^+^* (e) dots per *Th^+^* neuron in the CSMG of mice treated with vehicle (Veh.) or cerulein (Cer.) for 4 weeks. Each graph represents data pooled from three separate independent experiments for each receptor. Data on scattered dot plots are presented as mean ± SEM. Numbers of mice: vehicle, 3; cerulein, 3. Numbers of neurons analyzed: for *Gfra1* (b-c), vehicle, 721; cerulein, 716; for *Gfra2* (d-e), vehicle, 1051; cerulein, 781. Statistical analysis: Mann–Whitney test, with the corresponding *p*-values shown in the figure. (f) qPCR analysis of *Gfra1* or *Gfra2* mRNA expression in pancreatic FACS-sorted nm-pSCs from mice treated with vehicle (Veh.) or cerulein (Cer.) for 4 weeks. Data are presented as mean ± SEM. Each dot represents nm-pSCs coming from two pancreata and tested in triplicate for the vehicle or cerulein. Statistical analysis used the One sample Wilcoxon test, with the corresponding *p*-values shown in the figure. (g) Representative images of cultured sympathetic neurons isolated from the CSMG of mice treated with a vehicle or cerulein for 4 weeks. Immunostaining was performed using an anti-TH antibody. Neurons were plated in the presence of NGF with or without the addition of GDNF. Scale bar, 100 μm. (h) Scatter plots showing the mean axonal length of TH^+^ neurons from the CSMG of mice treated with vehicle (Veh.) or cerulein (Cer.) for 4 weeks. Data are presented as mean ± SEM. The graph represents the pool of three independent experiments using four mice per experiment. Numbers of neurons quantified: vehicle, 147 (NGF), 214 (NFG+GDNF); cerulein, 216 (NGF), 257 (NFG+GDNF). Statistical analysis: Kruskal–Wallis test with Dunn’s multiple comparisons, with the corresponding *p*-values shown in the figure. GDNF, glial cell-derived neurotrophic factor; CSMG, coeliac superior mesenteric ganglia; Th, Tyrosine Hydroxylase; Gfra1, GDNF Family Receptor Alpha 1; Gfra2, GDNF Family Receptor Alpha 2; nm-pSCs, non-myelinating pancreatic Schwann cells; NGF, nerve growth factor; Veh., vehicle; Cer., cerulein; w., week.

We then tested the effect of GDNF on cultured sympathetic neurons isolated from the CSMG. Adult sympathetic neurons showed very limited axonal extension *in vitro*, and, unlike embryonic axons (Ledda et al., 2002), they did not respond to the addition of GDNF to the culture medium (Figure 7g-h). In contrast, we observed an increase in axonal length in neurons from cerulein-treated mice compared with that in untreated controls (Figure 7g–h), suggesting that the intrinsic growth capacity of axons was stimulated by inflammatory conditions. In addition, the axonal outgrowth of cerulein-treated neurons was further enhanced in the presence of GDNF (Figure 7g–h). Thus, during CP, adult sympathetic neurons in the CSMG regained sensitivity to the axon growth-promoting activity of GDNF.

To address the relevance of GDNF release by nm-pSCs *in vivo*, we specifically depleted *Gdnf* in SCs by generating *Sox10-iCreER^T2^;Gdnf^Lox/Lox^* mice (Figure 8a). Under control conditions, the loss of *Gdnf* in SCs had no effect on the density of nm-pSCs or sympathetic nerve fibers in the exocrine pancreatic tissue (Figure 8b–c and Supp. Figure 7), indicating that GDNF expression was not necessary for the maintenance of neuroglial interactions under physiological conditions. However, after 2 weeks of cerulein treatment, the absence of GDNF abolished the increase in GFRA3^+^ surface density and GFRA3^+^ cell bodies normally observed in ADM lesions (Figures 8b and 8d–e). This effect was also associated with a striking reduction in the density of TH^+^ sympathetic nerve fibers in ADM lesions (Figures 8b and 8c). These data indicate that GDNF secretion by nm-pSCs was required to expand nm-pSCs in metaplastic pancreatic tissue and support sympathetic axon sprouting, possibly through a direct effect on GFRA1/2 expressing sympathetic neurons.

**FIGURE 8.**
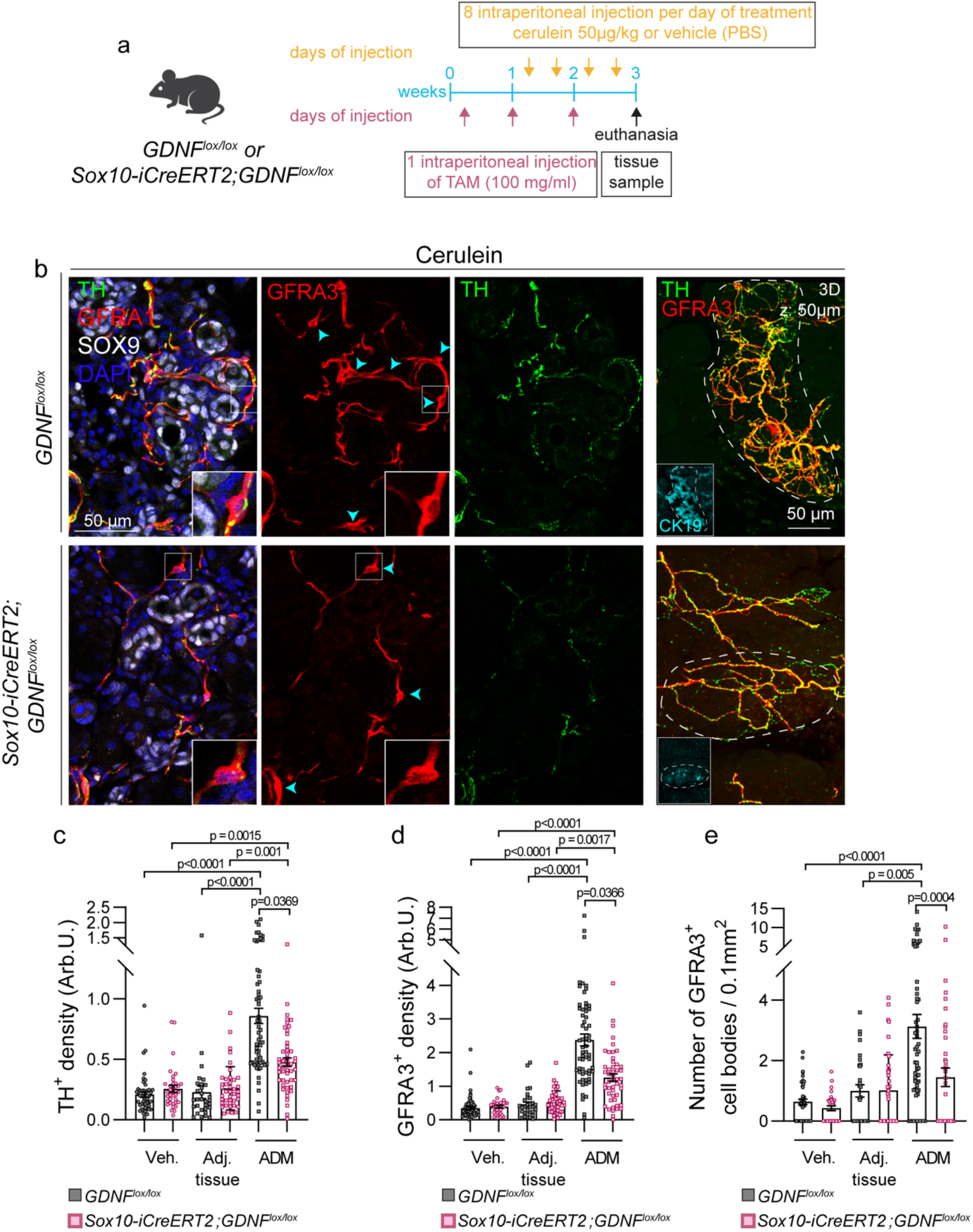
GDNF from nm-pSCs triggered SC expansion and axon sprouting. (a) Outline of the experimental procedure for combined genetic depletion of *Gdnf* in nm-pSCs and 3-week induction of chronic pancreatitis. (b) Representative images of cryostat sections and maximum intensity projection of 50 μm-thick optical sections through ADM regions in *Gdnf^Lox/Lox^* or *Sox10-iCreER^T2^;Gdnf^Lox/Lox^* mice immunostained with anti-GFRA3, anti-TH and anti-SOX9 or anti-CK19 antibodies. nm-pSCs cell bodies were pointed with cyan arrowheads and one nm-pSCs was magnified. Scale bars, 50 μm. (c–e) Scattered dot plots showing TH^+^ signal density (c), GFRA3^+^ signal density (d), and the number of GFRA3^+^ cell bodies (e) in the acinar parenchyma of mice injected with a vehicle (Veh.) or in the ADM and adjacent regions (adj. reg.) of mice injected with cerulein for 2 weeks. Data are presented as mean ± SEM. Statistical analysis: Kruskal–Wallis test with Dunn’s multiple comparisons, with the corresponding *p*-values shown in the figure. Numbers of mice: *Gdnf^Lox/Lox^*, 3 (vehicle) and 5 (cerulein); *Sox10-iCreER^T2^;Gdnf^Lox/Lox^*, 3 (vehicle), 4 (cerulein). Numbers of images analyzed for (c): *Gdnf^Lox/Lox^*, 48 (veh.), 30 (adj. tissue), 60 (ADM); *Sox10-iCreER^T2^;Gdnf^Lox/Lox^*, 36 (veh.), 41 (adj. tissue), 49 (ADM); for (d), *Gdnf^Lox/Lox^*, 65 (veh.), 30 (adj. tissue), 60 (ADM); *Sox10-iCreER^T2^;Gdnf^Lox/Lox^*= 36 (veh.), 41 (adj. tissue), 49 (ADM); for (e), *Gdnf^Lox/Lox^*, 52 (veh.), 29 (adj. tissue), 61 (ADM); *Sox10-iCreER^T2^;Gdnf^Lox/Lox^*, 36 (veh.), 41 (adj. tissue), 49 (ADM). GDNF, glial cell-derived neurotrophic factor; nm-pSCs, non-myelinating pancreatic Schwann cells; SC, Schwann cell; ADM, acinar-to-ductal metaplasia; SOX9, SRY-Box Transcription Factor 9; GFRA3, GDNF Family Receptor Alpha 3; TH, Tyrosine Hydroxylase; CK19, cytokeratin 19; veh., vehicle; adj., adjacent.

## 4 DISCUSSION

This study characterized the role of nm-pSCs in PDAC development. First, we showed that nm-pSCs changed in number and shape, enabling them to interact with new sympathetic branches that innervate the early precursor lesions of PDAC. Second, we found that nm-pSCs upregulated the expression of c-Jun, a master regulator of the repair phenotype, and the neurotrophic factor GDNF. Suppression of *Gdnf* in nm-pSCs inhibited sympathetic axon sprouting. These findings suggest that intra-organ SCs reprogramming supports neuronal remodeling during cancer initiation, which regulates tumor growth.

SCs have been implicated in promoting tumor growth and facilitating the metastatic dissemination of cancer cells through invading peripheral nerves (Deborde et al., 2022, 2016; Demir et al., 2014, 2017, 2016; Shurin et al., 2019; Sun et al., 2023; Weitz et al., 2023; Xue et al., 2023). These roles mainly result from the direct interactions between SCs and cancer cells and do not involve neurons. The present findings extend our current understanding of SC functions in tumorigenesis by revealing its involvement in the sprouting of sympathetic nerve terminals that innervate PDAC precursor lesions. The sympathetic nervous system plays a pro-tumorigenic role in certain cancers such as prostate and breast cancers (Kamiya et al., 2019; Magnon et al., 2013). However, selective sympathetic nerve ablation has shown opposite anti-tumorigenic functions in pancreatic cancer, in part by promoting an immunosuppressive macrophage phenotype (Guillot et al., 2022; Song et al., 2017). Therefore, the involvement of nm-pSCs in remodeling the sympathetic nervous system suggests that SCs may contribute to this protective anticancer activity. However, nm-pSCs have also been found to be associated with nonsympathetic fibers, including sensory fibers, which can promote pancreatic tumorigenesis (Saloman et al., 2016). Whether nm-pSCs also contribute to the remodeling and tumor infiltration of sensory fibers is likely but is yet to be demonstrated. Therefore, SCs may not be strictly pro-tumorigenic in cancer as previously though; rather, they may have dual functions depending on the type of cancer and the type of nerve fibers with which they interact.

Our findings support the hypothesis that nm-pSCs undergo adaptive reprogramming in response to chronic inflammation and early cancer development. However, the specific extrinsic signals that trigger this reprogramming remain unknown; these include the signals exchanged between neurons and SCs. For example, in nerve injury and neurodegenerative diseases, the loss of physical contact with axons and the release of “alarm” messengers, such as adenosine triphosphate, by degenerating axons triggers SC reprogramming (Negro et al., 2016; Rodella et al., 2017). The blockade of presynaptic neurotransmitter release induces axonal and terminal SC sprouting at the neuromuscular junctions (Son & Thompson, 1995). This observation supports the hypothesis that sympathetic nervous system activity, known to increase during inflammation, may contribute to the reprogramming of axon-attached nm-pSCs. Alternatively, the reprogramming of nm-pSCs can be triggered by the physical and chemical properties of the diseased pancreatic microenvironment. For example, increased tissue stiffness, as measured in an inflamed/premalignant pancreas (Payen et al., 2020; Rice et al., 2017), may affect SC morphology and c-Jun expression (Y. Gu et al., 2012; Rosso et al., 2022; Z. Xu, Orkwis, & Harris, 2021). SCs can be activated by the release of cytokines and growth factors, such as TGF-β (Clements et al., 2017; Fregnan, Muratori, Simões, Giuseppina, & Raimondo, 2012; Li et al., 2015; Shurin, Vats, Kruglov, Bunimovich, & Shurin, 2022; Sulaiman et al., 2002). These reprogramming signals originate from a variety of cells, most notably from premalignant/malignant cells. Conditioned media from cancer cells, including PDAC cells, can directly reprogram SCs (Deborde et al., 2016; Demir et al., 2014; Salvo, Saraithong, Curtin, Janal, & Ye, 2019; Shurin et al., 2019; Weiss et al., 2021; Zhou, Li, Han, Zhong, & Zhong, 2020), although the specific molecules involved remain unknown.

The current study demonstrated an expansion of the pancreatic SC network within lesions of the exocrine pancreas. Detailed analysis at single-cell resolution revealed remarkable morphological changes in nm-pSCs, characterized by the branching and extension of their processes. A previous study reported similar atypical morphologies of pancreatic SCs around islets in experimental diabetes models (Tang, Chiu, Hsu, Peng, & Fu, 2013), suggesting that nm-pSCs exhibit high structural plasticity, similar to other SC types. While a common c-Jun-dependent program may underlie SC plasticity, extrinsic signals and environmental cues are likely to fine-tune these programs, resulting in diverse morphologies tailored to perform specialized functions. For example, bipolar and very long repair SCs can bridge the gap between the two stumps of an injured nerve, providing a physical scaffold for axonal growth during regeneration (Gomez-Sanchez et al., 2017; Jessen & Arthur-Farraj, 2019). The complex star-like structure of the nm-pSCs observed here is likely crucial for providing a physical guide for the numerous branches that sprout from the sympathetic axon terminals. In healthy tissues, pronounced alignment between sympathetic axons and blood vessels suggests a reciprocal developmental interaction between these systems. In contrast, previous observations using the KIC model have shown minimal contact between sprouting sympathetic axons and blood vessels in pancreatic lesions (Guillot et al., 2022), suggesting that the pathological sprouting of sympathetic axons may not be significantly dependent on blood vessels. Instead, it is likely that the extended processes of the nm-pSCs provide a cellular substrate for the extension of collateral axons. However, further analyses are required to determine whether nm-pSCs extend these processes prior to axonal growth. Another, although not exclusive, possibility is that the branching and elongation of nm-pSCs serve to establish direct contact with cells in the local microenvironment. The present finding that nm-pSCs released inflammatory cytokines suggests their potential role in attracting immune cells to injury sites (Dubový, Klusáková, & Hradilová Svíženská, 2014; Napoli et al., 2012; Tofaris, Patterson, Jessen, & Mirsky, 2002). Therefore, the dynamic remodeling of nm-pSCs could help to position axon sprouts near immune cells, facilitating neuroimmune regulation in pancreatic diseases.

In addition to changes in the nm-pSC morphology, we observed an increase in their proliferation. SC proliferation also occurs after nerve injury but is not a critical factor for regenerative axon growth (Grinspan, Marchionni, Reeves, Coulaloglou, & Scherer, 1996; Kim et al., 2000; Yang et al., 2008). Instead, during development, this proliferation may be important for matching the number of SCs and nerve fibers. In pancreatic lesions, the morphological expansion of nonproliferating nm-pSCs may be insufficient to cover the increasing density of sympathetic axon sprouts after several weeks of cerulein treatment. This appears to require the generation of additional SCs, the number of which increased significantly after 4 weeks of treatment. The proliferative capacity of the nm-pSCs may also explain the recovery of normal cell numbers after conditional DTA-mediated ablation in the cerulein-induced CP model. A similar ability to regenerate the entire SC population at the neuromuscular junction and sciatic nerve following SC death was recently described (Gerber et al., 2019; Hastings, Mikesh, Lee, & Thompson, 2020).

Finally, we identified the role of GDNF in the function of nm-pSCs. GDNF is one of the best-known neurotrophic signals with increased expression in repair SCs (P. Xu et al., 2013). Application of exogenous GDNF or transplantation of engineered SCs overexpressing GDNF enhances nerve regeneration in models of peripheral nerve injury (Chen, Chai, Cao, Lu, & He, 2001; Hoyng et al., 2014; Santosa et al., 2013; Tannemaat et al., 2008). However, despite these findings, formal evidence that GDNF, similar to several other neurotrophic factors produced by repair SCs (including Shh, NGF, and BDNF, reviewed in Jessen & Mirsky, 2022, is required for nerve regeneration remains to be established. Here, we provide direct evidence that selective depletion of GDNF from SCs was sufficient to inhibit the sprouting of sympathetic nerve terminals in metaplastic pancreatic lesions. As sympathetic neurons and nm-pSCs express GFRA1 and GFRA2 receptors, GDNF can influence axonal remodeling by a direct action on axons, by an autocrine effect on nm-pSCs, or possibly by both mechanisms simultaneously Our in vitro experiments support a direct effect of GDNF on axons, which showed that adult sympathetic neurons from cerulein-treated mice exhibited increased axonal extension in the presence oledf recombinant GDNF. Signal transduction requires GFRA co-receptors in axons, such as RET (Proto-oncogene tyrosine-protein kinase receptor Ret) and the alternative transmembrane receptor NCAM (Durbec et al., 1996; Paratcha, Ledda, & Ibáñez, 2003; Trupp et al., 1996). Both receptors are expressed in developing sympathetic neurons (Enomoto et al., 2001; Mirsky, Jessen, Schachner, & Goridis, 1986) and, similar to the GFRA subunits, may be upregulated in adult sympathetic neurons that re-initiate axon growth.

GFRA1/NCAM signaling activates several intracellular pathways associated with SC migration and proliferation (Iwase, Jung, Bae, Zhang, & Soliven, 2005). GDNF induces SC proliferation in intact peripheral nerves when administered at high doses (Höke et al., 2003). Thus, autocrine GDNF signaling in nm-pSCs may be required for the observed expansion of pancreatic lesions, as evidenced by the reduced density of nm-pSCs in conditional *Gdnf* mice. This could indirectly affect the sprouting of sympathetic axon terminals, which would then lack the physical tracks to extend and possibly the energetic support provided by the SCs. Neuronal function is metabolically demanding, requiring continuous ion flow, transport of cellular cargo, and maintenance of a large membrane surface area. These requirements may be exacerbated during axonal growth. Given the considerable distance between the cell bodies of neurons and nerve terminals, it is unlikely that the intrinsic nutrient transport mechanisms of neurons are sufficient. The role of SCs in transporting metabolites, such as lactate, to axons is beginning to be characterized, and their importance in axonal regeneration after injury is emerging (Bouçanova & Chrast, 2020; Domènech-Estévez et al., 2015; Morrison et al., 2015). Thus, understanding how SCs respond to the metabolic and energetic demands of neuronal remodeling induced by cancer development is an interesting avenue for future research.

In conclusion, this study provides a novel and detailed characterization of a population of intra-organ SCs. Although pancreatic SCs may have organ-specific properties and functions, they share the capacity to proliferate and undergo reprogramming under pathological conditions with other SCs. This adaptability enables them to contribute to nervous system remodeling during chronic inflammation and early phases of tumorigenesis. The role of SCs in promoting the sprouting of uninjured axon terminals has been largely understudied but could potentially be a crucial and targetable process in diverse pathological conditions characterized by axonal sprouting, such as myocardial injury, chronic bone pain, diabetes, and joint and skin inflammation (Chartier et al., 2014; Kanazawa & Fukuda, 2022; Longo, Osikowicz, & Ribeiro-da-Silva, 2013). The findings of this study could provide clues for developing new therapies to modulate inflammation and cancer associated neural plasticity.

## ACKNOWLEDGMENTS

The authors would like to thank Oumaima Hattabi and Flavie Chung Shing for their substantial help with the analysis of Schwann cell morphology. We thank Pascal Durbec for providing insightful and constructive feedback on this manuscript. We thank Pascal Durbec, Xavier Bonnefont, Francoise Helmbacher, and Jaan-Olle Andressoo for providing the mouse strains used in this study. We also thank the Institute of Developmental Biology (IBDM) animal facility and France-BioImaging/PICsL infrastructure (ANR-10-INBS-04-01). This work was supported by the Centre National de la Recherche Scientifique (CNRS), France, Aix Marseille Université (AMU), France, and a grant from the Foundation Pour la Recherche Médicale (EQ202103012957) to FM. MMR-S received support from the French government under Programme Investissements d’Avenir, Initiative d’Excellence d’Aix-Marseille Université via A*Midex (AMX-19-IET-004) and ANR (ANR-17-EURE-0029) funding.

