## Supplementary data for "Pancreatic Schwann cell reprogramming supports cancer-associated neuronal remodeling"

### SUPLEMENTARY DATA

**Supp. Figure 1.** nm-pSCs associated with autonomic and sensory fibers in the mouse pancreas.

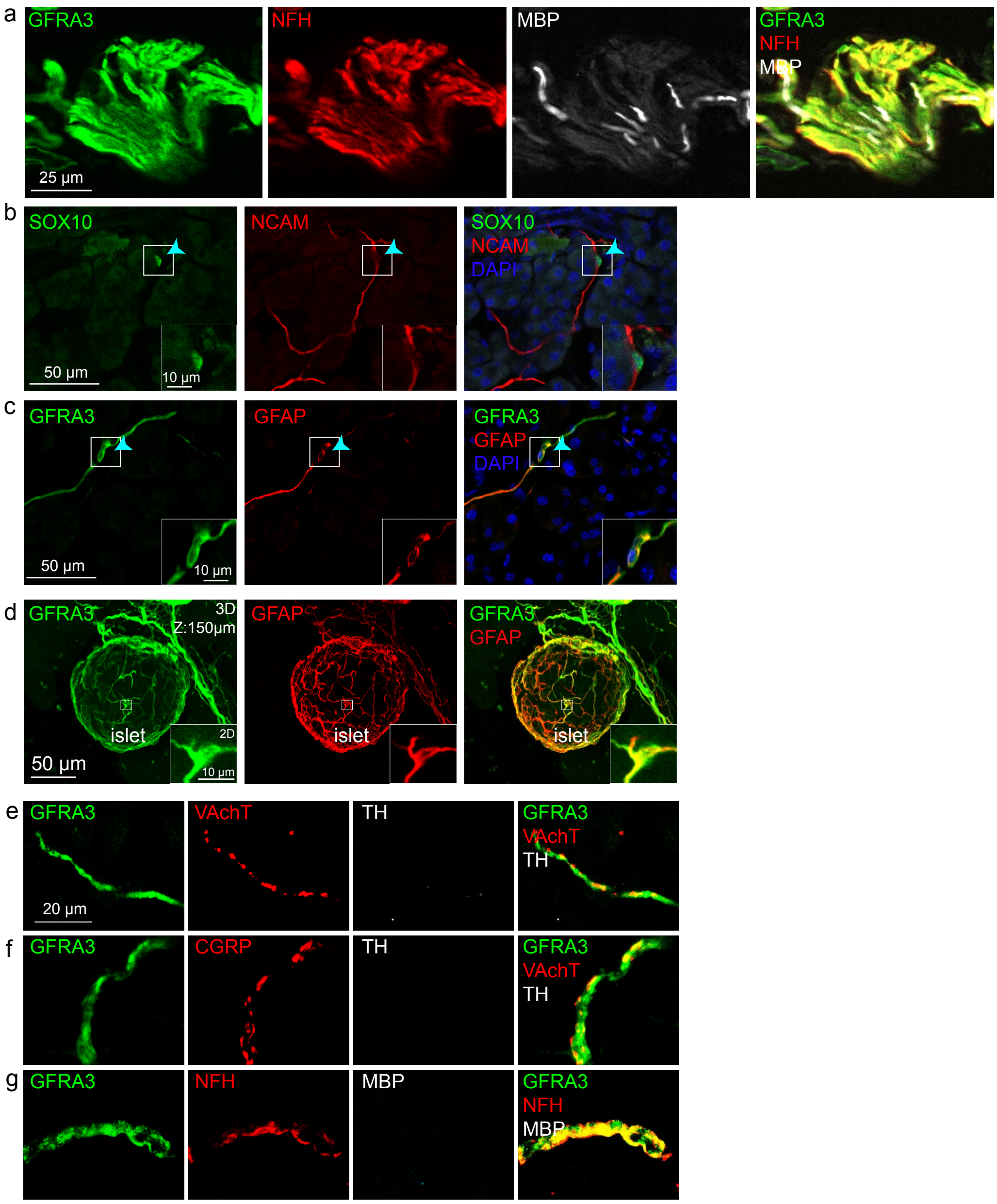


(a) Representative images of cryostat sections of an extrapancreatic nerve from an 8-week-old mouse immunostained for GFRA3, NFH, and MBP. Scale bar, 25 µm.

(b, c) Representative images of cryostat sections of mouse pancreas immunostained for GFRA3 and GFAP (b) or SOX10 and NCAM (c) to label nm-pSCs. The nm-pSCs was pointed with a cyan arrowhead and magnified for better observation. Scale bar, 50 µm.

(d) Maximum intensity projection of a 150 μm-thick optical section through a cleared pancreas showing staining with anti-GFRA3 and anti-TH antibodies in an islet of Langerhans. One nm-pSC cell body was magnified. Scale bar, 50 μm.
(e–f) Representative images of cryostat sections of mouse pancreas immunostained for GFRA3, TH, VACHT, CGRP, NFH, and MBP. Scale bar, 20 μm.
All images are representative observations in at least three mice.
nm-pSCs, non-myelinating pancreatic Schwann cells; GFRA3, GDNF Family Receptor Alpha 3; NFH, neurofilament, heavy polypeptide; MBP, myelin basic protein; GFAP, glial fibrillary acidic protein; SOX10, SRY-Box Transcription Factor 10; NCAM, neural cell adhesion molecule; TH, Tyrosine Hydroxylase; VACHT, vesicle choline acetyltransferase; CGRP, calcitonin gene-related peptide.

**Supp. Figure 2.** nm-pSCs associated with autonomic and sensory fibers in KIC pancreas.

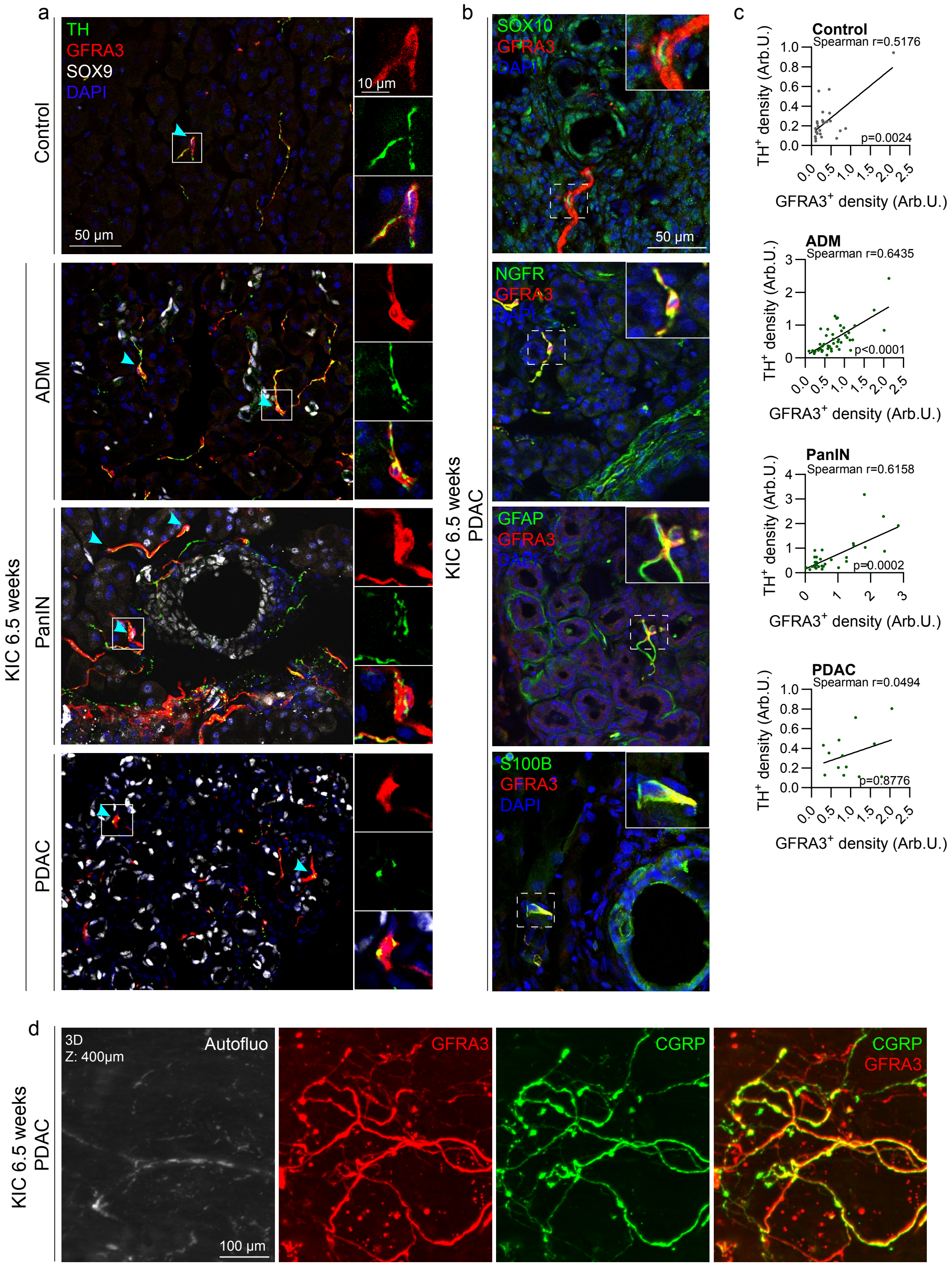

(a) Representative images of pancreatic sections immunostained with anti-TH, anti-GFRA3, and anti-SOX9 antibodies showing the acinar parenchyma of a control mouse and ADM, PanIN, or PDAC lesions of a 6.5-week-old KIC mouse. nm-pSCs cell bodies were pointed with cyan arrowheads. In the insets, the cell body of the nm-pSCs was magnified to illustrate their close proximity to the $TH^+$ fibers. Images are representative of those observed in pancreatic samples from at least three mice. Scale bars, 50 µm and 10 µm in the magnified panel.
(b) Representative images of cryostat sections of a 6.5-week-old KIC PDAC tumor immunostained for GFRA3, SOX10, NGFR, GFAP, and S100B. The cell nuclei were counterstained with DAPI. In each inset, one cell body of the nm-pSCs was magnified. Images are representative examples of three examined mice. Scale bars: 50 µm in the magnified panel.
(c) Correlation analysis between densities of $TH^+$ fibers and $GFRA3^+$ nm-pSCs within healthy pancreas, ADM, PanIN and PDAC from KIC mice. Statistical analysis was performed using the Spearman test, the corresponding *p*-values and correlation coefficient are shown in the figure.
(d) Maximum intensity projection of a 400-µm-thick optical section through the PDAC of a 6.5-week-old KIC mouse showing immunolabeling with anti-GFRA3 and anti-CGRP antibodies and autofluorescence (Autofluo) of the tissue. The images are representative of pancreatic samples from the three pancreases. Scale bar, 100 µm.

nm-pSCs, non-myelinating pancreatic Schwann cells; KIC, $\textit{Kras}^{\textit{LSL}\text{-G12D/+}}$; $\textit{Cdkn2a (Ink4a/Arf)}^{\textit{lox/lox}}$*; Pdx1-Cre*; TH, Tyrosine Hydroxylase; GFRA3, GDNF Family Receptor Alpha 3; SOX9, SRY-Box Transcription Factor 9; ADM, acinar-to-ductal metaplasia; PanIN, pancreatic intraepithelial neoplasia; PDAC, pancreatic ductal adenocarcinoma; SOX10, SRY-Box Transcription Factor 10; NGFR, nerve growth factor receptor; GFAP, glial fibrillary acidic protein; CGRP, calcitonin gene-related peptide.

**Supp. Figure 3.** Coverage of sprouting sympathetic fibers by nm-pSCs in chronic pancreatitis.

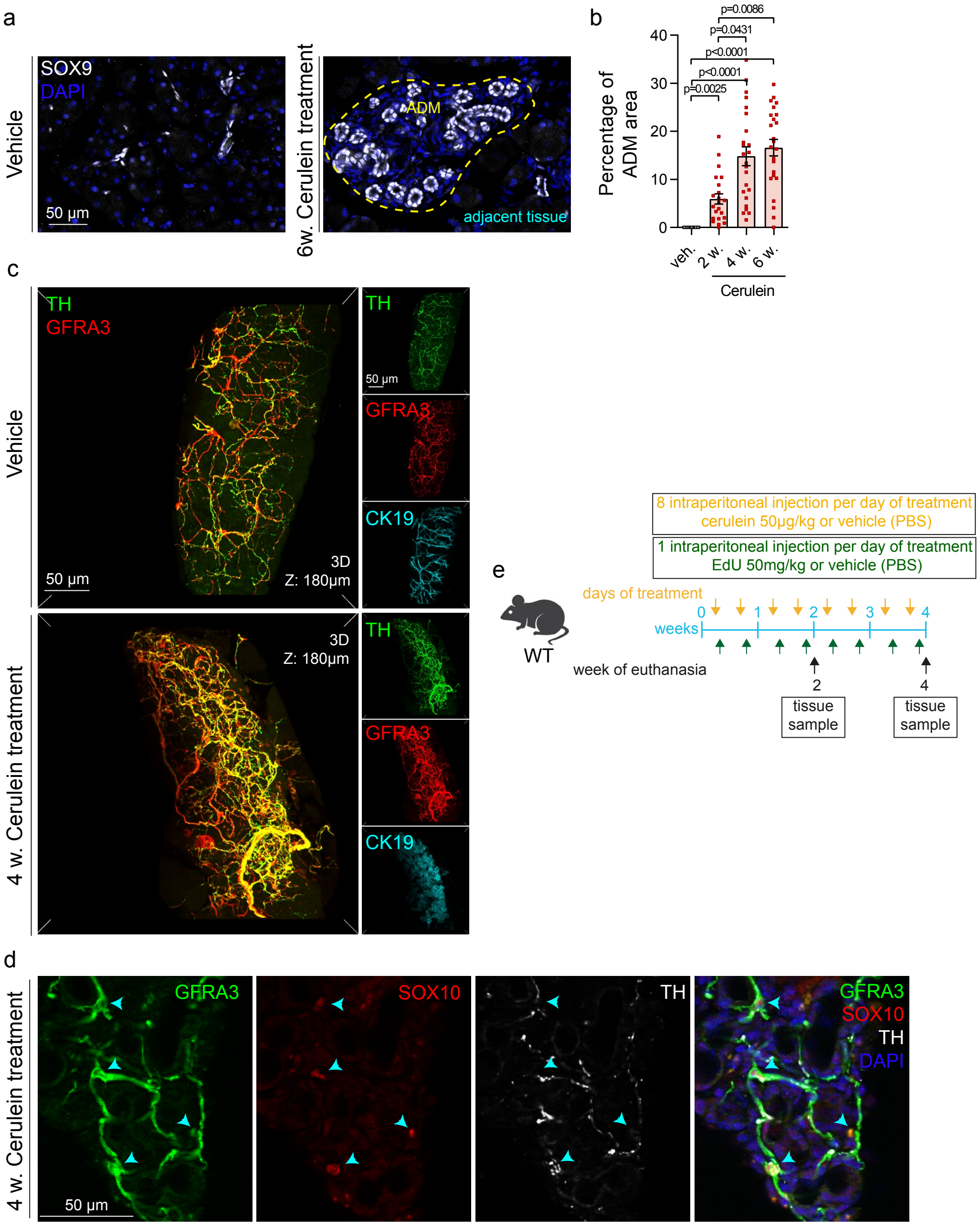


(a) Representative images of pancreatic cryostat sections immunostained with anti-SOX9 antibody showing the acinar parenchyma in vehicle-treated mice (a) or ADM lesions in mice treated with cerulein for 4 weeks (b). The cell nuclei were counterstained with DAPI. Scale bars, 50 μm.

(b) Scattered dot plots showing the percentage of the ADM areas in pancreatic sections from mice injected with vehicle (veh.) or cerulein for 2, 4, or 6 weeks (w). Data are presented as mean ± SEM. Statistical analysis: Kruskal–Wallis test with Dunn's multiple comparisons, with corresponding *p*-values shown in the figure. Numbers of mice: vehicle, 3; cerulein, 4. Numbers of images analyzed: vehicle, 20; cerulein, 22 (2 weeks), 24 (4 weeks), and 24 (6 weeks).

(c) Maximum intensity projection of 180-μm-thick optical sections through cleared pancreas from mice injected with vehicle or cerulein for 4 weeks showing triple labeling with anti-GFRA3, anti-TH, and anti-CK19 antibodies. A scaled-down version of individual staining was displayed to the right of each panel. Images are representative of metaplastic lesions observed in the pancreas of more than three mice. Scale bars, 50 μm and 50 μm in panels with individual staining.

(f) Representative images of pancreatic cryostat sections immunostained with anti-GFRA3, anti-SOX10, and anti-TH antibodies in ADM lesions induced by cerulein injection after 4 weeks of treatment. The cell nuclei were counterstained with DAPI. nm-pSCs cell bodies were pointed with cyan arrowheads. Images are representative of metaplastic lesions observed in the pancreas of more than three mice. Scale bars, 50 μm.

(g) Outline of the CP experiment combined with EdU injections.

nm-pSCs, non-myelinating pancreatic Schwann cells; SOX9, SRY-Box Transcription Factor 9; ADM, acinar-to-ductal metaplasia; GFRA3, GDNF Family Receptor Alpha 3; TH, Tyrosine Hydroxylase; CK19, cytokeratin 19; CP, chronic pancreatitis; veh., vehicle; w., week.

**Supp. Figure 4.** Validation of nm-pSC reporter mice.

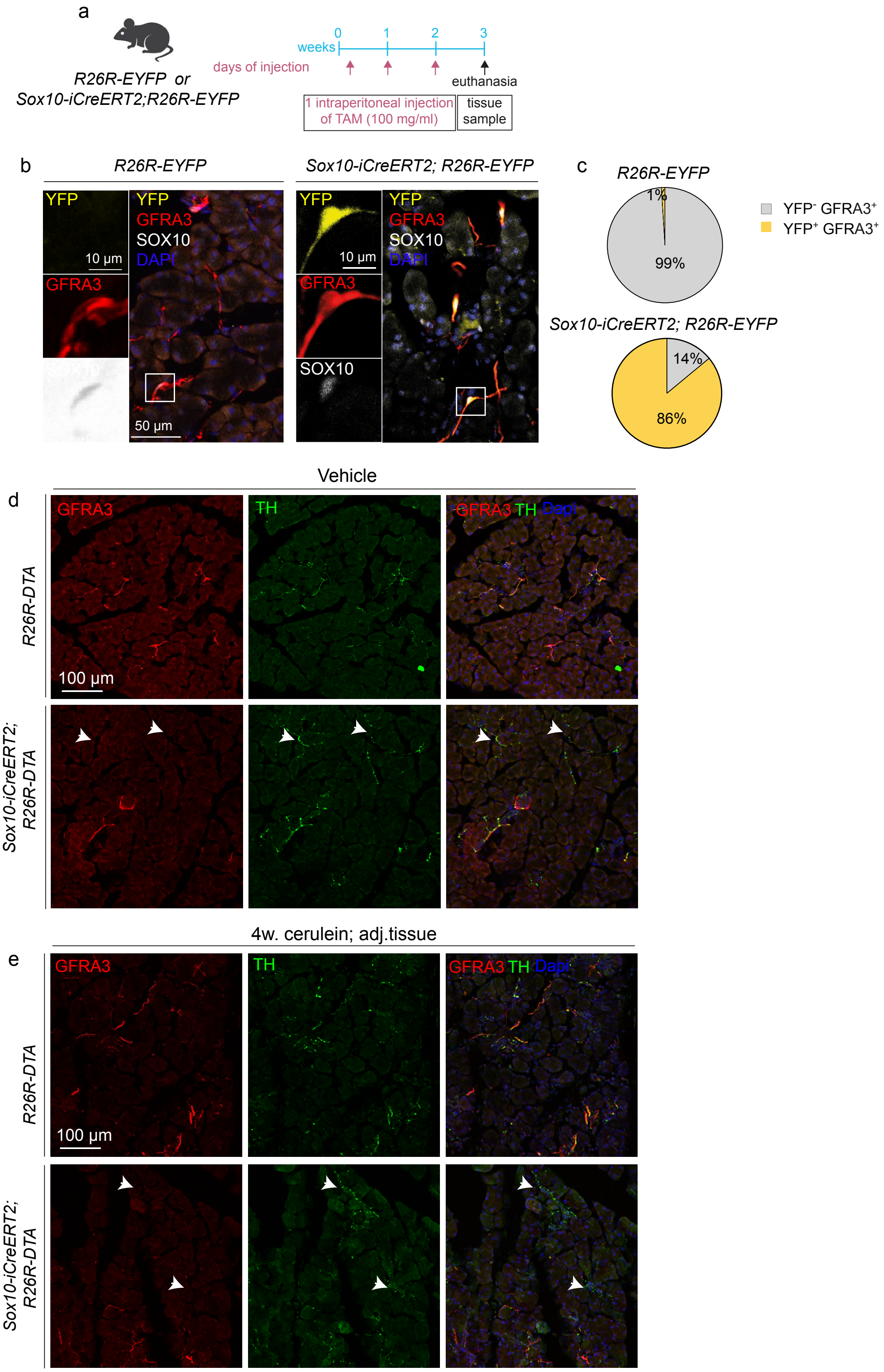

(a) Outline of the experimental procedure for genetic YFP staining of nm-pSCs from *Sox10-iCreERT2;R26R-YFP* mice.

(b) Representative images of pancreatic cryostat sections from *R26R-YFP* and *Sox10-iCreER$^{T2}$;R26R-YFP* mice immunostained with anti-YFP, anti-GFRA3, and anti-SOX10. The cell nuclei were counterstained with DAPI. In each panel, one cell body of the nm-pSCs was magnified. Scale bars, 50 µm and 10 µm in the magnified panel.

(c) Pie charts showing the percentages of YFP$^-$GFRA3$^+$, YFP$^+$GFRA3$^+$, and YFP$^-$GFRA3$^-$ cells in the pancreas illustrated in (b). *R26R-YFP*: n=3 mice; analyzed cells, 78 (77 YFP$^-$GFRA3$^+$, 1 YFP$^+$GFRA3$^+$). *Sox10-iCreERT2;R26R-YFP*: n=3 mice; analyzed cells, 122 (17 YFP$^-$GFRA3$^+$, 105 YFP$^+$GFRA3$^-$).

(d) Representative images of pancreatic sections from vehicle-injected *R26R-DTA* and *Sox10-iCreER$^{T2}$;R26R-DTA* mice immunostained for anti-GFRA3 and anti-TH antibodies. The cell nuclei were counterstained with DAPI. TH$^+$ fibers without nm-pSC were found after SC depletion (white arrowheads). Images are representative of findings observed in at least three mice. Scale bar, 100 µm.

(e) Representative images of pancreatic cryostat sections from *R26R-DTA* or *Sox10-iCreERT2;R26R-DTA* mice treated with cerulein for 2 weeks showing immunostaining for anti-GFRA3 and anti-TH antibodies in histologically asymptomatic adjacent tissues. TH$^+$ fibers without nm-pSC were found after SC depletion (white arrowheads). Images are representative of findings observed in at least three mice. Scale bars, 100 µm.

nm-pSCs, non-myelinating pancreatic Schwann cells; YFP, yellow fluorescent protein; GFRA3, GDNF Family Receptor Alpha 3; SOX10, SRY-Box Transcription Factor 10; TH, Tyrosine Hydroxylase.

**Supp. Figure 5.** Custom-made 3D-printed imaging chambers and morphology analysis of nm-pSCs.

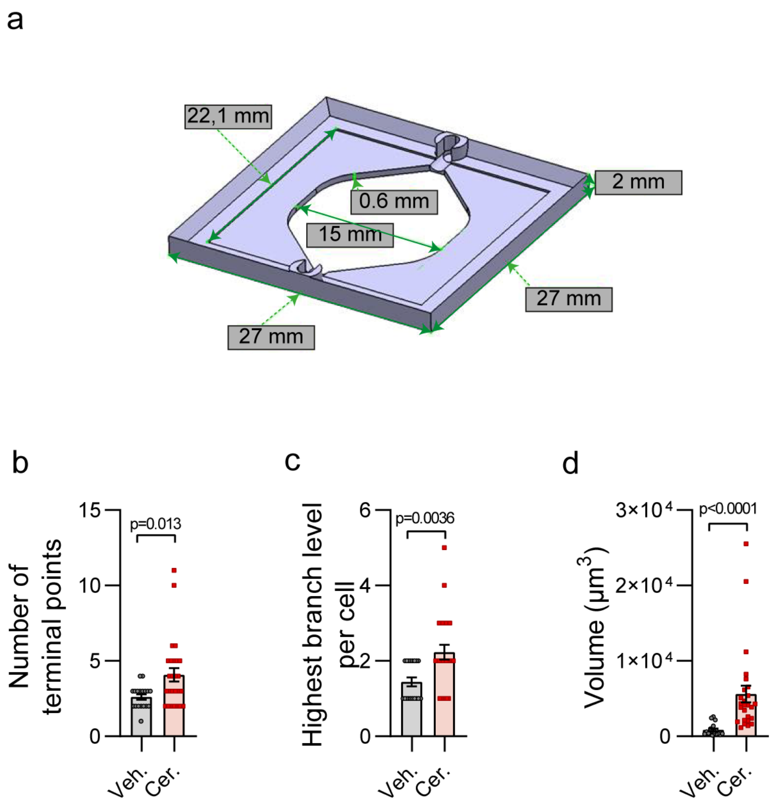


(a) Design of 3D-printed chambers for imaging-cleared 500-µm-thick pancreatic slices.

(b–d) Quantification of the number of terminal points (b), highest branch level (c), and volume (d) of nm-pSCs in the pancreas of mice treated with vehicle (Veh.) and cerulein (Cer.) for 2 weeks. Data are presented as mean ± SEM. Vehicle: n=3 mice, 18 nm pSCs; cerulein: n=3 mice, 26 nm pSCs. Statistical analysis was performed using the Mann-Whitney test, and the corresponding *p*-values are shown in the figure.

nm-pSCs, non-myelinating pancreatic Schwann cells. Veh., vehicle; Cer., cerulein.

**Supp. Figure 6.** FACS sorting of nm-pSCs.

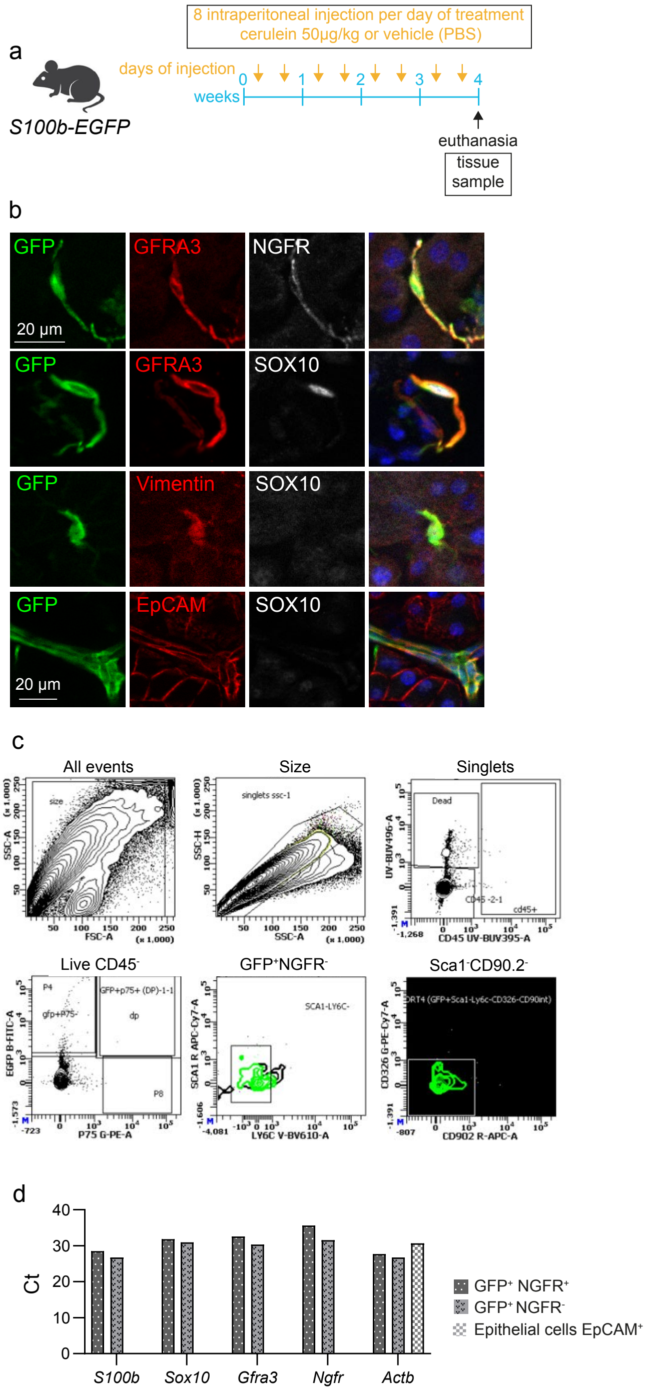


(a) Outline of induced-chronic pancreatitis experiment in *S100b-EGFP* mice.

(b) Representative images of cryostat sections of the pancreas from *S100b-EGFP* mice showing GFP expression in GFRA3$^{+}$, NGFR$^{+}$ and SOX10$^{+}$ nm-pSCs, EpCAM$^{+}$ epithelial cells, and vimentin$^{+}$ fibroblasts Images are representative observations in at least three mice. Scale bars, 20 µm.

(c) FACS gating strategy for isolating nm-pSCs from *S100b-EGFP* mouse pancreata. The cells were gated according to their sizes and granularity, and doublets were subsequently removed. Live/CD45$^{-}$ cells were gated to remove dead cells and immune cells; within this population, GFP+NGFR+ and GFP+NGFR- cells containing nm-pSCs were gated. Finally, to remove epithelial cells and fibroblasts from the GFP+ cells, Ly6C/Sca1-/CD326/CD90.2 cells were gated, and the resulting cells were sorted as SCs.

(d) qPCR analysis of mRNA expression of *S100b, Sox10, Gfra3, Ngfr, Actb* in FACS-sorted GFP$^{+}$ NGFR$^{+}$, GFP$^{+}$ NGFR$^{-}$ nm-pSCs, and GFP$^{+}$ EpCAM$^{+}$ pancreatic epithelial cells from vehicle-treated mice.

nm-pSCs, non-myelinating pancreatic Schwann cells; GFP, green fluorescent protein; GFRA3, GDNF Family Receptor Alpha 3; NGFR, nerve growth factor receptor; SOX10, SRY-Box Transcription Factor 10; EpCAM, Epithelial cell adhesion molecule; S100b, S100 Calcium Binding Protein B; Actb, actin beta; Ct: cycle threshold.

**Supp. Figure 7.** GDNF deletion from nm-pSCs had no effect on SC expansion and axon sprouting in control pancreata and the regions adjacent to ADM.

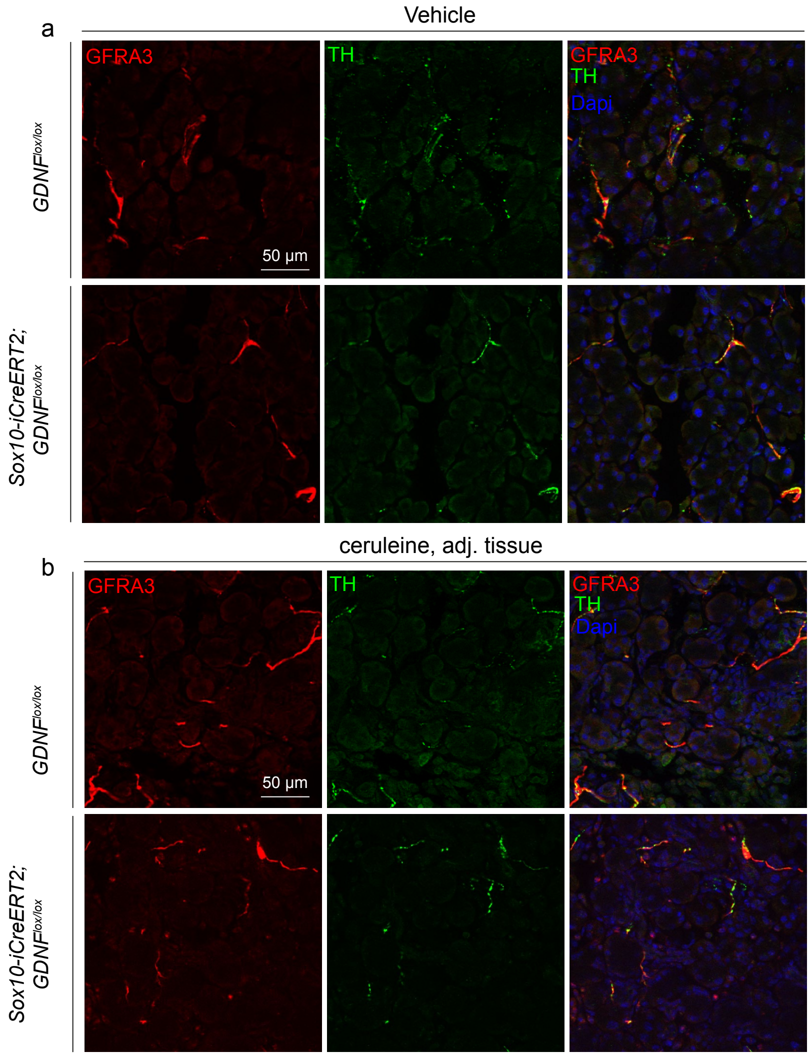


(a) Representative images of pancreatic cryostat sections from vehicle-treated *GDNF*$^{lox/lox}$ and *Sox10-iCreER*$^{T2}$*; GDNF*$^{lox/lox}$ mice, immunostained with anti-GFRA3 and anti-TH antibodies. Images are representative of findings observed in at least three mice. Scale bar =50 μm.

(b) Representative images of pancreatic cryostat sections from *GDNF*$^{lox/lox}$ and *Sox10-iCreER*$^{T2}$*; GDNF*$^{lox/lox}$ mice treated with cerulein for 4 weeks showing anti-GFRA3 and anti-TH antibody staining in the histologically asymptomatic tissues adjacent to the ADM (adjacent region). Images represent metaplastic lesions observed in the pancreas of more than three mice. Scale bar =50 μm.

GDNF, glial cell-derived neurotrophic factor; SC, Schwann cell; ADM, acinar-to-ductal metaplasia; GFRA3, GDNF Family Receptor Alpha 3; TH, Tyrosine Hydroxylase.
